# Spatiotemporal dynamics of human microglia are linked with brain developmental processes across the lifespan

**DOI:** 10.1101/2021.08.07.455365

**Authors:** D.A. Menassa, T.A.O. Muntslag, M. Martin-Estebané, L. Barry-Carroll, M.A. Chapman, I. Adorjan, T. Tyler, B. Turnbull, M.J.J. Rose-Zerilli, J.A.R. Nicoll, Z. Krsnik, I. Kostovic, D. Gomez-Nicola

## Abstract

Microglia, the brain’s resident macrophages, shape neural development and wiring, and are key neuroimmune hubs in the pathological signature of neurodevelopmental disorders. In the human brain, microglial development has not been carefully examined yet, and most of our knowledge derives from rodents. We established an unprecedented collection of 97 postmortem tissues enabling quantitative, sex-matched, detailed analysis of microglial across the human lifespan. We identify the dynamics of these cells in the human telencephalon, describing novel waves in microglial density across gestation and infancy, controlled by a balance of proliferation and apoptosis, which track key neurodevelopmental milestones. These profound changes in microglia are also observed in bulk RNAseq and single-cell RNAseq datasets. This study provides unparalleled insight and detail into the spatiotemporal dynamics of microglia across the human lifespan. Our findings serve as a solid foundation for elucidating how microglia contribute to shaping neurodevelopment in humans.

## INTRODUCTION

Microglia are the main resident immune cells of the brain. During development, their roles include phagocytosis of neuronal precursors to restrict progenitor pool size (Cunningham et al., 2013); the modulation of forebrain wiring by guiding interneuron positioning (Squarzoni et al., 2014); the pruning of synapses (Paolicelli et al., 2011); and the formation and refinement of axonal tracts (Pont-Lezica et al., 2014; Squarzoni et al., 2014; Verney et al., 2010). In the adult, microglia retain key homeostatic functions, suppressing interneuron activation and modifying animal behaviour (Badimon et al., 2020); regulate hippocampal neurogenesis (Sierra et al., 2010); and drive inflammation in a disease context (Gomez-Nicola and Perry, 2015).

However, most of our knowledge of microglial developmental dynamics is derived from rodent studies. In the mouse, microglia originate from erythromyeloid progenitors in the extraembryonic yolk sac (YS) from E7.5 (Ginhoux et al., 2010). These progenitors begin populating the brain primordium at E9.5 and their phenotypic specification into microglia is defined by intrinsic and brain-specific transcriptional regulators (Bennett et al., 2018; Gosselin et al., 2014; Lavin et al., 2014). The entire microglial population is generated by expansion during embryonic life and the early postnatal developmental stages, followed by a transient postnatal selection phase (Askew et al., 2017; Réu et al., 2017). Under homeostatic conditions, the population is maintained throughout life by cycles of slow selfrenewal estimated at approximately 0.69% per day (Askew et al., 2017).

Human microglial development has some similarities to mouse, as macrophages are detectable by 2-3 postconceptional weeks (pcw) in the blood islands of the extraembryonic YS (Janossy et al., 1986; Nogales, 1993; Park et al., 2020; Popescu et al., 2019), to later appear in the forebrain from the 3^rd^ pcw (Menassa and Gomez-Nicola, 2018; Verney et al., 2010). A series of descriptive post-mortem studies collectively report on microglial proliferation in clusters appearing in the ventral telencephalon and diencephalon from the 4^th^ pcw and continuing throughout fetal development (Monier et al., 2007; Monier et al., 2006; Verney et al., 2010). Recent single-cell transcriptomic studies suggest that microglial ontogenic pathways are conserved between human and mouse during embryonic development (Bian et al., 2020), with microglia progressing from an undifferentiated state towards a mature adult-like immunocompetent state from 11 pcw in humans (Kracht et al., 2020). The early arrival of microglia to the brain precedes the onset of pivotal processes of human cortical development such as neurogenesis, neuronal migration and gliogenesis (Menassa and Gomez-Nicola, 2018). In the adult, the population self-renews at a slow daily turnover rate which has been estimated at 0.08-2% (Askew et al., 2017; Réu et al., 2017). However, compared with the mouse, we lack a deep understanding of microglial spatiotemporal dynamics during human development, largely due to the scarcity of developing tissues available for research.

We established an unprecedented collection of 97 post-mortem tissues enabling quantitative, sex-matched, and detailed analysis of microglial dynamics across the human lifespan (3^rd^ pcw – 75 years old), in relation to their immunocompetent and neurogenetic roles. We identify novel developmental and postnatal waves of expansion and refinement, where the population undergoes marked changes in numbers, and we determine the contributions of proliferation, apoptosis and migration to this process. We validate our identified critical windows in further analyses of datasets from 251 bulk RNAseq samples and 4 single-cell RNAseq (scRNAseq) studies of 24,751 microglial cells spanning the embryonic and fetal ages (3-24 pcw). This study is pivotal for our understanding of microglial biology and serves as a solid basis for elucidating how microglia shape the neurodevelopmental landscape in humans.

## RESULTS

### Brain colonisation by microglia begins on the 4^th^ postconceptional week (CS12)

The onset of embryonic circulation is concomitant with the development of the cardiovascular system, which in humans becomes functional from the late 3^rd^/early 4^th^ pcw (somites: 4-12; CS10 onwards) (O’Rahilly and Muller, 2010; Tavian and Peault, 2005). At CS10, we identify early colonisation of embryonic organs such as the heart by cells committed to the myeloid lineage (PU.1^+^; Fig 2A), with expression of IBA1 not yet detected (Fig 2A). By CS12 (the end of the 4^th^ pcw), organ colonisation by IBA1^+^ cells (Fig 2B) is profuse, with liver macrophage proliferation peaking at CS16 and accounting for the increase in macrophage cell densities observed from CS18 to CS21 (Fig 2C).

**Figure 1.**
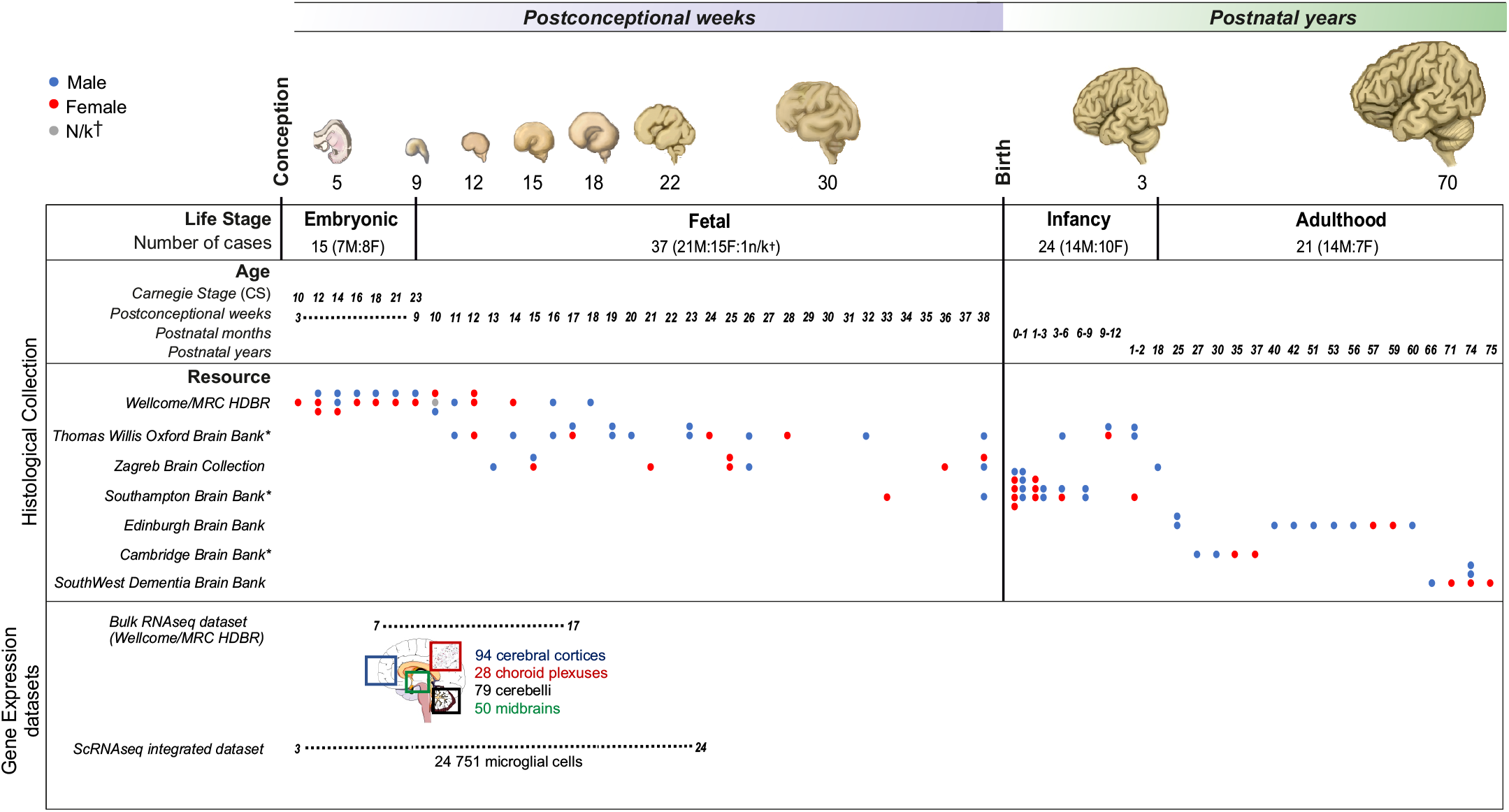
Overview of the study. (A) Post-mortem collection of human tissues across the lifespan with 52 prenatal and 45 postnatal cases which adds up to a total of n = 97. Gene expression datasets: bulk RNAseq of 251 samples from the Wellcome MRC/HDBR resource in 4 anatomical regions between 7-17 postconceptional weeks and an integrated dataset of 24751 microglial cells from 4 singlecell RNAseq studies (Bian et al., 2020; Cao et al., 2020; Fan et al., 2020; Kracht et al., 2020) between 3-24 postconceptional weeks. *F*: Female; *M*: Male; *MRC/HDBR*: Medical Research Council/Human Developmental Biology Resource; *†*Not known; *\**Brain banks that are part of the BRAINUK network.

**Figure 2.**
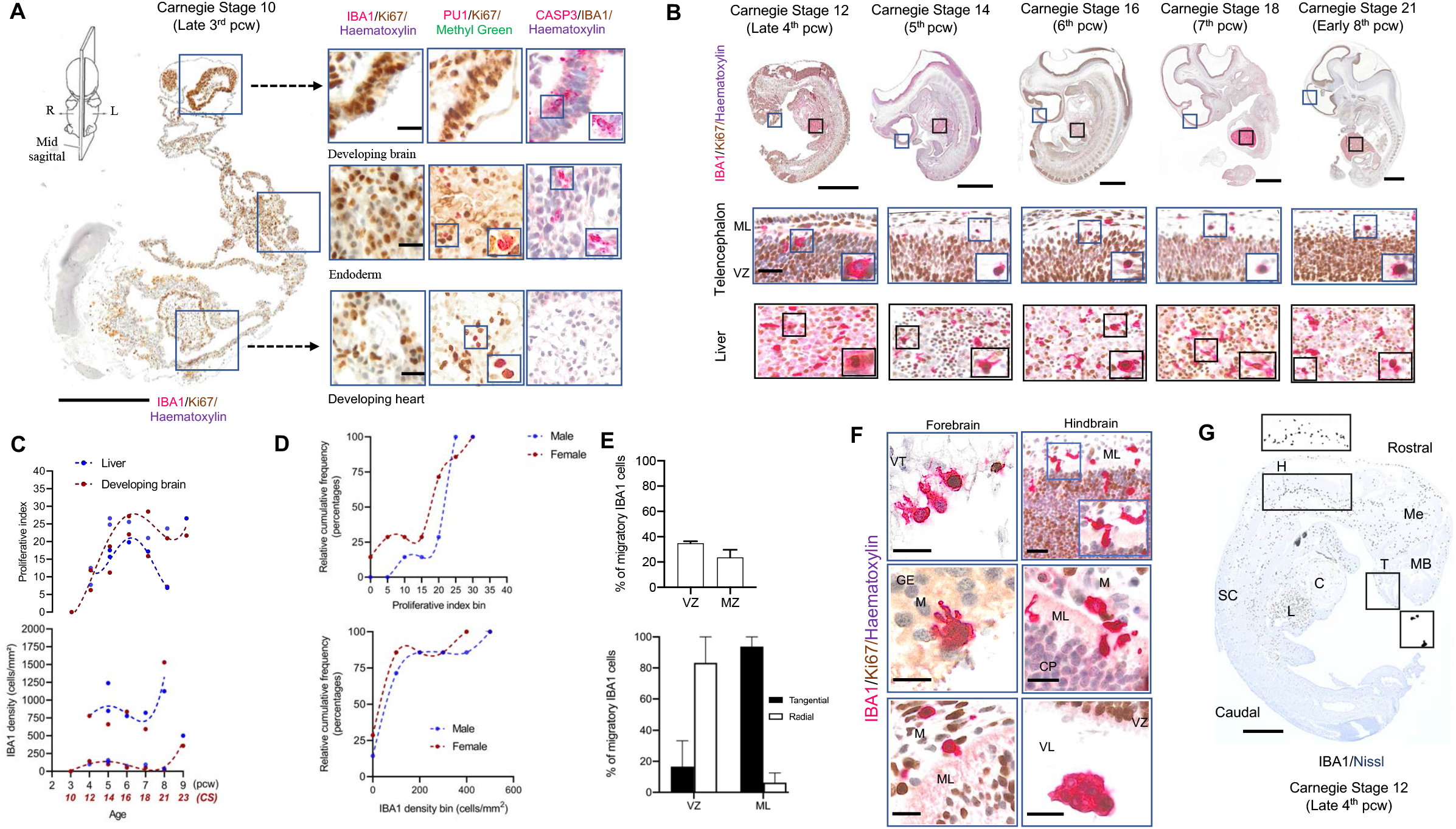
Early colonisation of the human embryo. (A) Representative sagittal cross-section through a CS10 (late 3^rd^ pcw) human embryo. Immunolabeling was done on consecutive 8 um sections and identifies the presence of PU.1 cells in organs apart from the brain and the absence of IBA1 cells overall. Scale bars: 500 μm (left); 40 μm (right). (B) Representative sagittal sections through whole human embryos from CS12, the age of appearance of IBA1 cells across the entire embryo, through to CS21. Insets show the location of proliferating IBA1 cells in both the liver and the dorsal telencephalon. Scale bars: 1 mm (top); 100 μm (bottom). (C) IBA1 proliferation dynamics and cell densities in the developing brain (n=15) and the liver (n=11). Lines represent a fitted loess function followed by a smoothing spline function for visualisation. Liver densities and proliferation are significantly higher compared to the developing brain (Wilcoxon test, p<0.001). For the brain, densities increase significantly from the 9^th^ pcw compared to earlier timepoints (Mann-Whitney U, p=0.01). (D) Cumulative distributions of densities and proliferative indices by sex in the developing brain (7F:7M) (2-sample Kolmogorov-Smirnov test; p>0.05). E) Representative migratory profiles of IBA1 cells in the bilaminar telencephalon at 5 pcw (n=2). (F) Entry routes of IBA1 cells into the brain rudiment. Scale bar: 100 μm. (G) IBA1 cell distributions in the CS12 embryo. Scale bar: 500 μm. *C*: cardiac muscle; *CP*: cortical plate; *GE*: ganglionic eminence; *H*: hindbrain; *L*: liver; *M*: meninges; *MB*: midbrain; *Me*: mesenchyme; *ML*: mantle layer; *SC*: spinal cord; *T*: telencephalon; *VZ*: ventricular zone.

However, at CS10, no IBA1^+^ or PU.1^+^ myeloid cells are detected in the brain rudiment (Fig 2A). By CS12, the first wave of brain colonisation begins, with IBA1^+^ macrophages seeding the forebrain, midbrain and hindbrain (Fig 2B). Microglial densities and proliferation vary between regions: cell densities are highest in the hindbrain at CS12 (Fig 2G) through to CS21 (Supplementary fig 7A) and proliferation is highest in the midbrain (Supplementary fig 7A). Microglial proliferation peaks at CS16 accounting for the first expansion phase of the population with a sharp increase in IBA1^+^ cell density from CS23 (Fig 2C). We found no significant differences between the cumulative distributions of microglia by sex for both density and proliferation in the developing embryonic brain (Fig 2D). Early in development (5^th^ pcw), migratory microglia (IBA1^+^) account for ~40% of the total population in the dorsal telencephalon, which at this age is bilaminar with a proliferative zone, the VZ, and a plexiform mantle layer (ML) (Fig 2E, top). Most of migration in the VZ is radial, indicating the colonization of adjacent layers, whereas microglia in the ML are characterised by a tangential migratory phenotype, suggesting expansion within a layer (Fig 2E, bottom).

In sum, we can date the arrival of microglial progenitors to the brain at CS12, with the population rapidly engaging into proliferative and migratory activities.

### The density of the microglial population fluctuates by waves of proliferation followed by cell death in the developing cortex

The future neocortex is formed from the dorsal telencephalon in the forebrain, which is where we focused all our subsequent analyses. Between CS12 and CS21, microglial proliferation is restricted to the ML (Fig 2B) and cells are seen entering the brain parenchyma from the VZ, the meningeal compartment and the GE (Fig 2F).

The last stage of embryonic life is CS23 (O’Rahilly and Muller, 2010), equivalent to the late 8^th^/early 9^th^ pcw, followed by fetal life. We observed two key proliferation waves characterising the expansion of the microglial population, both during fetal life:

*The first wave* is the most significant and coincides with the appearance of the cortical plate (Fig 3A) at the transition between embryonic and fetal life (late 8^th^/early 9^th^ pcw) (Kostovic et al., 2002; O’Rahilly and Muller, 2010). This increase in microglial proliferation (IBA1^+^Ki67^+^) predicts the peak in density observed subsequently at 10-12 pcw (Fig 3B-C) in the telencephalon, which is now characterised by 6 transient layers (Fig 3A, supplementary fig 2A). After 12 pcw, the microglial density drops to baseline levels (Fig 3B) due to microglial apoptosis that we detect by cleaved Caspase 3 labelling in microglia between 12-14 pcw (Fig 3D). Between 9-10pcw we also detect a significant transient switch in the migratory phenotype of microglia, with most cells adopting a radial migratory pattern (Fig 3E), across all telencephalic layers (Supplementary Fig 8), suggestive of a local colonisation process being coupled to cycles of expansion.

**Figure 3.**
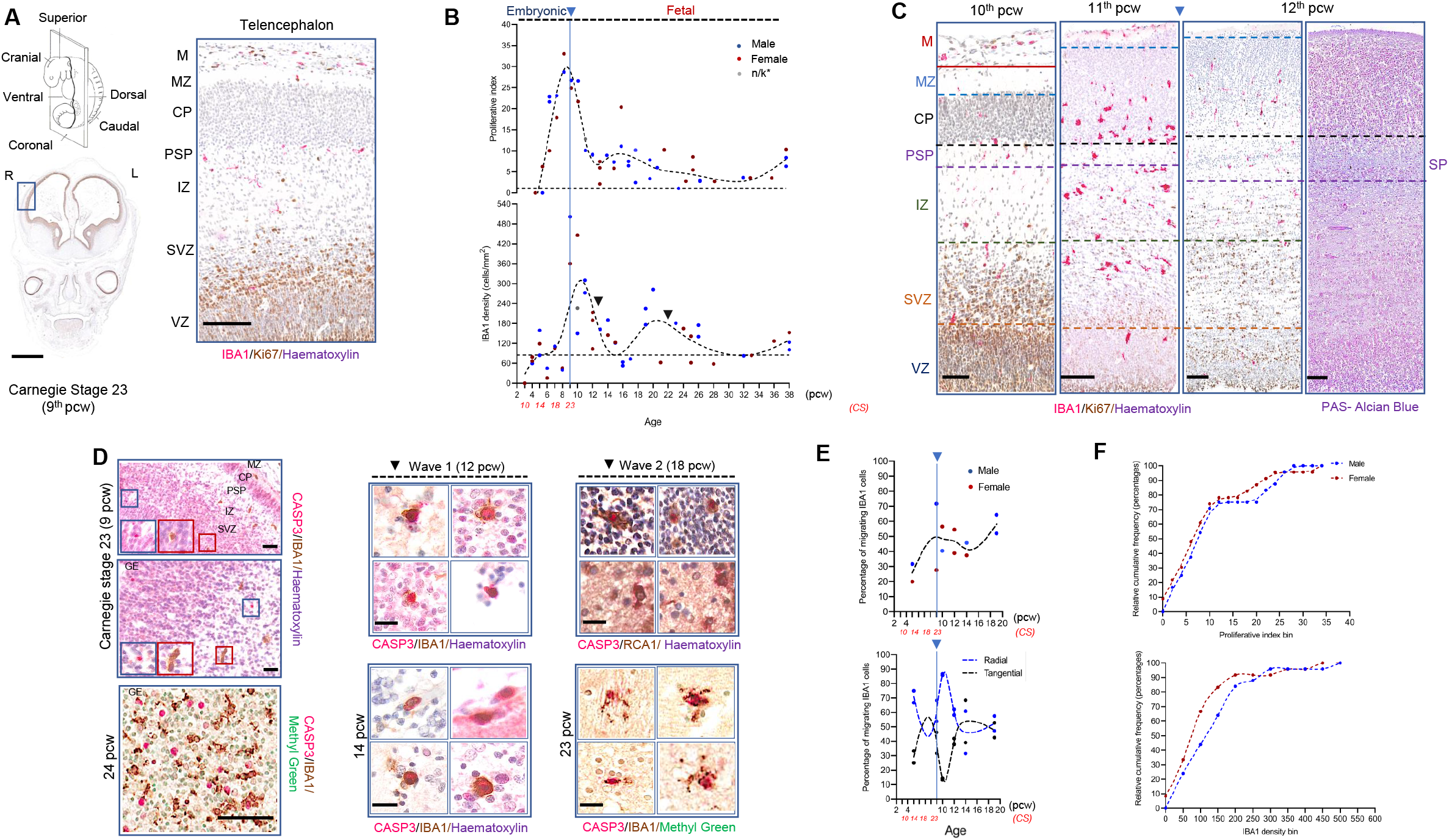
Developmental dynamics of microglia in the cortex. (A) Representative laminar structure of the developing cortex with its transient zones observed from CS23 (9^th^ pcw) in humans. Scale bar: 2mm (left); 100 μm (right). (B) Corrected microglial densities (against fold change in telencephalic thickness with age) and proliferative dynamics in the telencephalon during development (CS10 (late 3^rd^/early 4^th^ pcw) until term (38 pcw) (n = 50). \**n/k*: not known. Lines represent a fitted loess function followed by a smoothing spline function for visualisation. The most significant proliferation peak is between 5-9 pcw compared to 3-4 pcw (Kruskal-Wallis test, p<0.0001),10-21 pcw (Kruskal-Wallis test, p=0.0063) and 23-38 pcw (Kruskal-Wallis test, p<0.0001). Significant density peaks were between 9-15 pcw compared with 3-8 pcw (Kruskal-Wallis test, p<0.0001) and 16-32 pcw, the second peak (Kruskal-Wallis test, p=0.02) (C) Representative cortical columns from 10-12 pcw showing 1) the development of the pre-subplate to the subplate at 12.5 pcw and the alignment of microglial cells at the CP-PSP/SP boundary and 2) the distribution of microglial cells across transient zones. Scale bar: 100 μm. (D) Non-microglial cell death in the GE and cortical transient zones at CS23 (9^th^ pcw) and at 24 pcw (Left; scale bar: 100 μm); microglial cell death observed in two stages following the decrease in densities (black arrows in (B,D)) (Right; scale bar: 10 μm). (E) Migratory profile of microglia and type of migration in representative cases from the telencephalon (n = 12). (F) Proliferative dynamics and densities by sex across human development (2-sample Kolmogorov-Smirnov test, p>0.05). *R*: right; *L*: left; *CP.* cortical plate; *GE*: ganglionic eminence; *IZ*: intermediate zone; *M*: meninges; *MZ*: marginal zone; *PSP*: pre-subplate; *SP*: subplate; SVZ: subventricular zone; *VZ*: ventricular zone.

*The second wave* of proliferation coincides with the early expansion of the human subplate between 13-16 pcw (Kostovic, 2020) (Fig 3B). Proliferation predicts the increase in density peaking at 20 pcw (Fig 3B). Thereafter, the density drops again to baseline levels and closely matches the mean values obtained from adult densities (mean N_D_ = 84.63± 4.41 mean cells/mm^2^±SEM). This drop in density can be explained by microglial apoptosis (cleaved Caspase 3^+^) between 18-24 pcw (Fig 3D).

The wave-like pattern followed by the microglial population during development is consistently observed in both females and males (Fig 3B, 3E, 3F), with a trend towards an earlier onset of the proliferative waves observed in females (Fig 3F).

The microglial expansion is not uniform across transient zones of the developing cortex. As development progresses, we observed a pattern of “hot spots” of local proliferation preceding an increase in local density in subsequent timepoints, which is replicated across layers (Fig 4A-D). Every layer has a different timing: whereas densities peak earlier in development for the MZ, CP and SP (Fig. 4A-C), they do so much later for the SVZ and the VZ (Fig 4A, 4D). Migration patterns vary within each layer (Supplementary fig 8) perhaps indicating movement of cells between adjacent layers.

**Figure 4.**
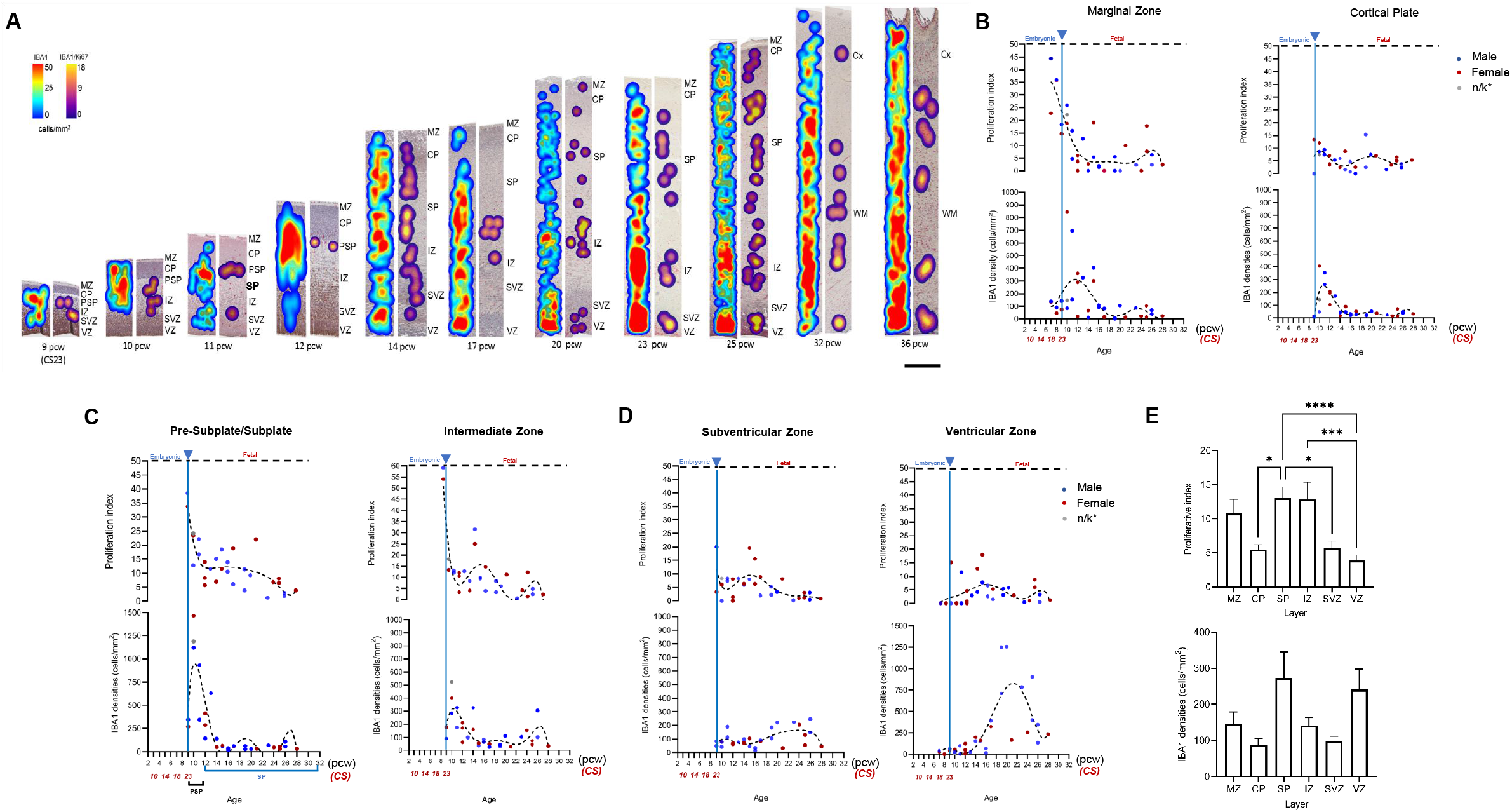
Spatiotemporal microglial dynamics within the developing cortex. (A) Representative heatmaps showing microglial dynamics throughout development in the human cortex. Heatmaps were constructed based on the density of immunoreactive cells. Density of all IBA1 immunoreactive cells are represented by blue-red spectral heatmaps (left column of each stage) while IBA1+/Ki67 double-positive cells are represented by purpleyellow spectral heatmaps (right column of each stage). Approximate densities are shown on the colour bar with minimal density being blue/purple, maximum density being red/yellow. All cortical columns are 450 μm wide. (B) Proliferation and densities of microglial in the marginal zone (n=35), cortical plate (n=31); (C) the subplate (n=31) and the intermediate zone (n=31); (D) the subventricular (n=31) and the ventricular zones (n=35). (E) Proliferation and densities in the subplate and the intermediate zones (Friedman’s test, p< 0.001). Lines represent a fitted loess function followed by a smoothing spline function for visualisation. *CP*: cortical plate*; GM;* grey matter; *IZ*: intermediate zone*; MZ*: marginal zone; *PSP*: presubplate; *SP*: subplate; *SVZ*: subventricular zone; *VZ*: ventricular zone: *WM*: white matter.

Altogether, post-colonisation at CS12, the microglial population follows a wave-like pattern of significant increases in density caused by proliferation, later refined by cell death, with the ultimate consequence of densities stabilising at levels observed in the adult after every wave.

### Microglia undergo transcriptional changes that mirror the wave-like pattern of the population dynamics

We sought to validate our histological findings with an alternative approach based on the analysis of the transcriptional profile of microglia. Using bulk RNA-seq data from whole brain (Gerrelli et al., 2015; Lindsay et al., 2016), we examined the expression of genes characteristic of the adult microglial signature (Supplementary file 2) across development (7-17 pcw). We considered a gene to be constitutively expressed when present in >80% of the samples in every timepoint at a TPM value >2 (Fig 5A). We identified that 24% (212) of cortical genes from the adult signature were constitutively expressed in the telencephalon between 8-17 pcw (ON genes; Fig 5A). These genes are involved in the regulation of the innate immune and inflammatory responses as well as cytokine production (Fig 5A). Heatmap representation of a set (75) of highly enriched microglial genes (Supplementary file 2) indicated a highly changeable signature with more expression in the earlier timepoints (8 and 9 pcw) compared to later timepoints whereby only a few of genes had high expression by 13-17 pcw (Fig 5B). The homeostatic marker P2RY12 had a stable expression throughout, whilst CX3CR1 increased gradually towards the 13-17 pcw temporal window (Fig 5C). Pro-inflammatory factors such as IRF5 and IFNGR1 had a stable expression throughout. The sensome gene CD37 had a 100-fold higher expression compared to other genes in our samples, declining with age. AIF1, encoding the ubiquitously expressed IBA1 protein, was expressed at low levels but consistently across the various ages (Fig 5C).

**Figure 5.**
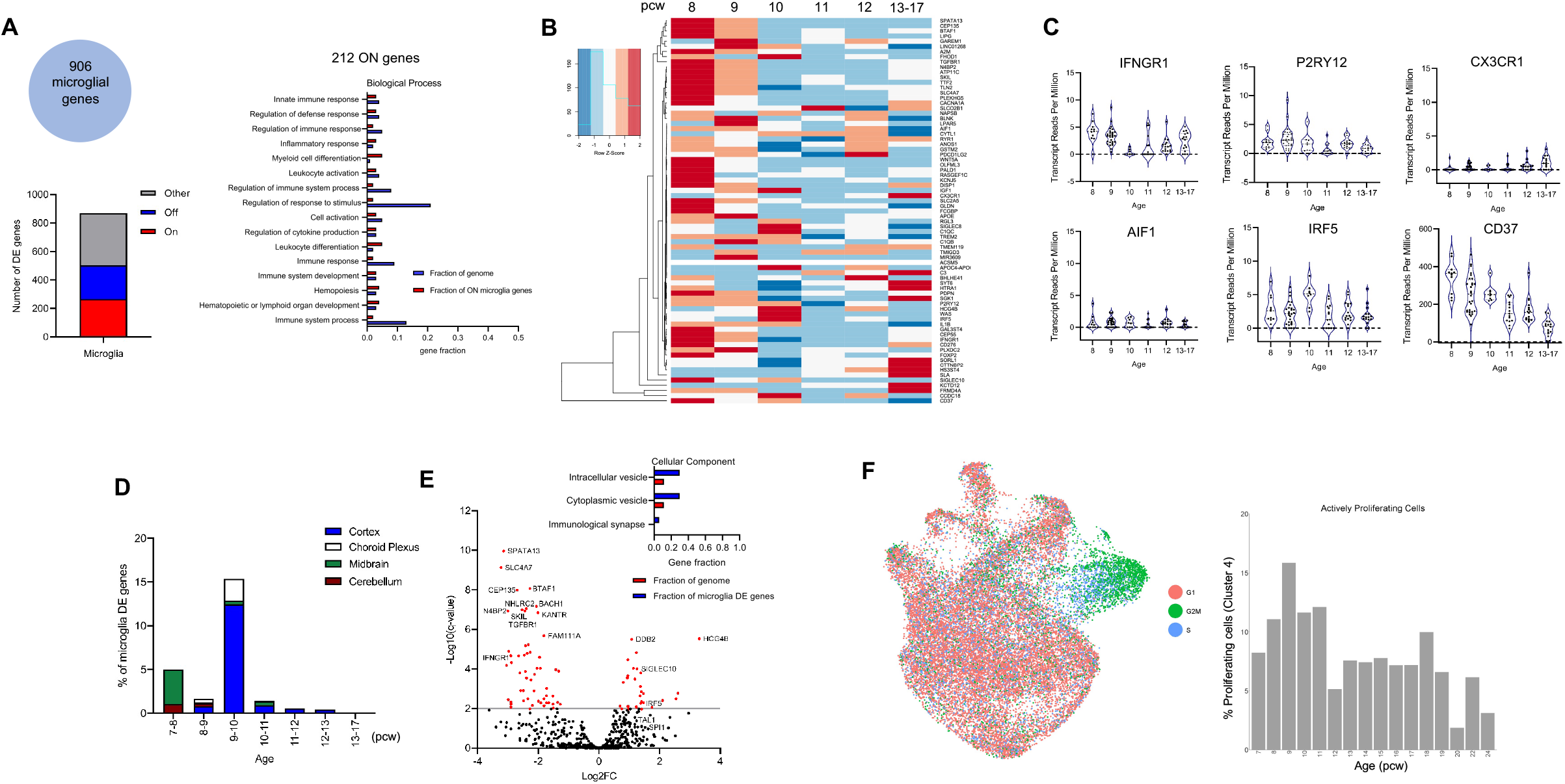
Microglial gene signature during development. (A) 906 genes chosen from two published microglial cortical adult and juvenile gene lists from (Galatro et al., 2017; Gosselin et al., 2017). Bar plot shows the percentage of constitutively expressed microglial genes, unexpressed genes and those with sporadic expression in the developing cortex with gene ontology analysis of constitutively expressed genes only. (B) Heatmap of highly enriched microglial genes during development (n = 75). (C) Transcript reads per million expression levels of a selection of homeostatic and immune modulatory microglial markers during development. (D) Regional microglial signatures based on cortical adult microglial genes. (E) Volcano plot of upregulated and downregulated microglial genes at the peak of differential gene expression at 9-10 pcw with gene ontology analysis significant for cellular component. (F) Actively cycling and proliferating microglia (CPM) in the scRNAseq dataset were identified across gestational age. Only ages with a minimum of 50 cells were selected (7 – 24 PCW). CPM signatures in single-cell data display a bimodal pattern of proliferation consistent with histological findings. *DE*: differentially expressed genes.

We compared the expression of the microglial signature by region, testing samples from the cerebellum, the choroid plexus, the cortex and the midbrain (Fig. 5D). We detected a significant increase in the %DE genes in the midbrain and cerebellum at 7-8 pcw, not seen in the cortex and choroid plexus until 9 pcw, suggesting different regions follow different maturation timings (Fig 5D). Thereafter, change in gene expression by region declined substantially with age (Fig 5D).

The peak of DEG detected at 9 pcw in the cortex matched the timing of the first wave of expansion of the microglial density (Fig 3). At 9pcw, we detected 81 microglial DEG: 27 upregulated DEG associated with chemokine and toll-like receptor signalling such as IRF5, SIGLECs (e.g. SIGLEC10) and MHC-II/co-stimulatory molecules such as HCG4B (Fig 5E & supplementary file 2) and 54 downregulated DEG associated with cytokine-cytokine receptor signalling and interferon-gamma mediated signalling such as TGFBR1 (Fig 5E). Collectively, microglial DEG are significantly associated with cellular components of immunological synapses and cytoplasmic, as well as, intracellular vesicles (Fig 5E). A further analysis of an integrated human scRNA-seq myeloid dataset from 4 sources (Bian et al., 2020; Cao et al., 2020; Fan et al., 2020; Kracht et al., 2020) identified 10 clusters of microglia (Supplementary Fig 6) with a distinct cluster (cluster 4) of actively cycling and proliferating microglial cells. This proliferation cluster also displayed a wave-like distribution across ages, first peaking at 9 pcw and then peaking again at 18pcw (Fig 5F, Supplementary Fig 6), tracking the pattern observed at the histological level.

### Microglial proliferation is prominent in key neurodevelopmental structures

The fetal subplate (SP) is a major site for synaptogenesis, neuronal maturation and a waiting compartment for cortical afferents (Kostovic, 2020). Microglia are enriched in the PSP monolayer below the CP from the 10^th^ pcw (Fig 3C, Fig 4A). The SP is fully formed at 12.5 pcw by the gradual merging of the PSP and the deep loose portion of the CP in parallel with the development of the afferent fibre system (Kostovic, 2020). Microglia appear to track these key changes, appearing densely clustered in the SP with microglial proliferation highest in the SP and the IZ (Fig 4E, Supplementary fig 9A-F).

By 15-20 pcw, we observed significant clustering of microglia in periventricular crossroads of projection and callosal pathways in the frontal lobe; these cells co-express MHCII and are non-proliferative (Supplementary Fig 9). By 20 pcw, microglia are evenly distributed throughout the expanding SP and only penetrate the upper portion of the CP by 25 pcw (Supplementary Fig 9B). Dense clusters of microglia can be observed in the VZ from 22-28 pcw, colonising all the cortical layers by 28 pcw (Supplementary Fig 9B).

From 32 pcw, the 6-layered cortical Grundtypus can be observed with a resolving SP zone. Grey (GM) and white matters (WM) are clearly identifiable and regional differences between GM and WM can already be measured (Supplementary Fig 10). Both density and proliferation decreased significantly and level out with adult densities (Supplementary Fig 10).

In sum, we found qualitative evidence of the intimate association of microglia with the key neurodevelopmental hallmarks that characterise brain formation and wiring, suggestive of a bidirectional crosstalk with neuronal progenitors.

### The microglial population undergoes an expansion phase during infancy and is refined to adult levels by cell death

Postnatally, the microglial density increases from birth and, around 1-2 years of age, we observe a *third wave* characterised by a 4-fold increase in comparison with the adult and third trimester densities (Fig 6A). This increase is predicted by a rise in proliferation just after birth and until 6 months of age (Fig. 6A). Thereafter, proliferation decreases to the levels observed in the adult, remaining constant through life (Fig 6A). The mechanism through which microglial densities decrease after 1.5 years may be likened to the selection phase observed in rodents postnatally (Askew et al., 2017; Nikodemova et al., 2015) and we indeed detect microglial death (RCA1^+^Caspase3^+^) in the human cortex (Fig 6B-C). We found no significant differences between the cumulative distributions by sex for both density and proliferation (Fig 6D). The adult density and proliferation of microglia retain the previously described (Mittelbronn et al., 2001) higher density in the white matter (Fig 6E, 6F, 6G). From 18 years of age, throughout adult life and healthy ageing, the population has a slow turnover of 1.12 + 0.18 % and the density stabilises at 84.63 ± 4.42 cells/mm^2^ (Fig. 6G).

**Figure 6.**
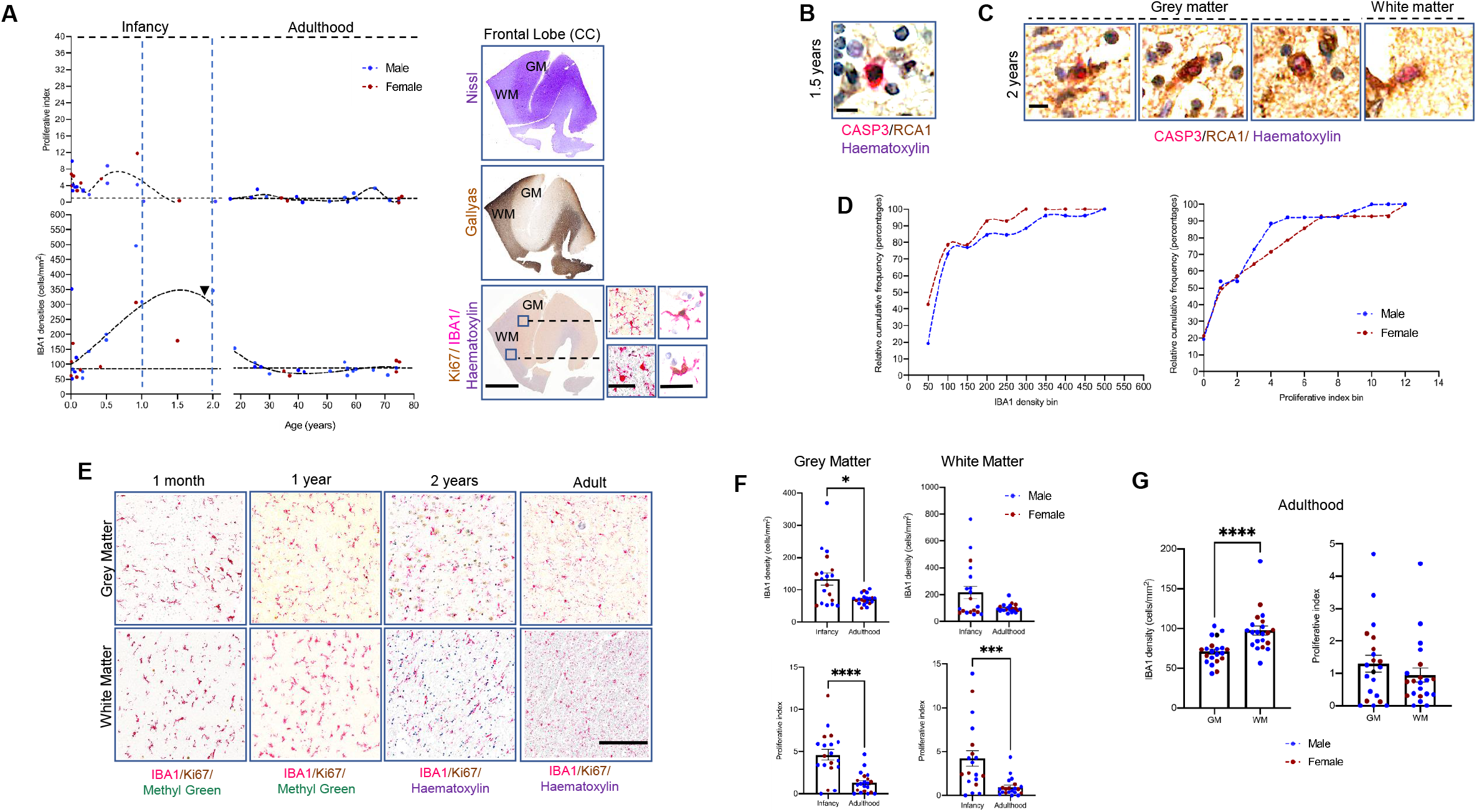
Postnatal dynamics of microglia in the cortex. (A) Microglial densities and proliferative dynamics in infancy (n =24; 14M:10F) and adulthood (n = 21; 14M:7F) and a representative cross-section of frontal cortex with anatomical histochemistry and photomicrographs of homeostatic and proliferating microglial morphologies in grey and white matters. Lines represent a fitted loess function followed by a smoothing spline function for visualisation. Infancy proliferation peak was most significantly different between 0.4-2 years compared with birth-0.4 years (Mann-Whitney U, p=0.0015). Densities were significantly highest between 0.5-2 years compared to earlier timepoints (Mann-Whitney U, p<0.0001). Comparing infancy (0-2 years) with the adult (18-75 years) rates for proliferation, it is significantly highest in infancy (Mann-Whitney U, p<0.0001). This is similar for the density which is highest in infancy (0-2 years) compared to adult stages (Mann-Whitney U, p=0.04). Scale bar: 2 mm (left); 40 μm (right). (B) Representative photomicrograph of cell death in a non-microglial cell at 1.5 years. Scale bar: 15 μm. (C) Microglial cell death in grey and white matter at 2 years (black arrow in (A)). Scale bar: 10 μm. (D) Cumulative distributions of microglial densities and proliferation by sex. (2-sample Kolmogorov-Smirnov test; p = 0.09). (E) Representative photomicrographs of cell densities in grey and white matters from birth to 2 years. Scale bar: 500 μm. (F) Regional differences of microglial densities and proliferation in grey and white matters between adulthood and infancy. (G) Proliferation rates and densities in adulthood in grey and white matters*. CC: cingulate cortex; GM*: grey matter; *WM*: white matter.

## DISCUSSION

The past decade saw an exponential increase in experimental studies focusing on microglia, in part fuelled by the resolution of a long debate on the origins of these cells in mouse, by fate-mapping studies (Ginhoux et al., 2010). In humans, microglia enter the brain primordium prior to the onset of substantial neurogenesis and neuronal migration (Menassa and Gomez-Nicola, 2018). Recent studies indicate that microglia show marked heterogeneity during development, becoming immunocompetent from the 10^th^ pcw (Kracht et al., 2020) with conserved ontogenic pathways between mouse and human (Bian et al., 2020). The population expands during embryonic life to yield the adult population, which maintains itself by cycles of self-renewal at a slow daily turnover rate (Askew et al., 2017; Nikodemova et al., 2015; Réu et al., 2017). Due to limited availability of healthy developmental human tissues, most of our knowledge of microglial development stems from rodent studies. Such studies have highlighted the roles that microglia play in shaping the neurodevelopmental landscape, including neuronal genesis, migration, dendritic development, axonal pruning, synaptogenesis and wiring (Paolicelli et al., 2011; Squarzoni et al., 2014; Verney et al., 2010). However, until now we lacked a reference map for the development of microglia in humans, as an essential framework to contextualise basic studies and target human-relevant mechanisms. Here, we have characterised the precise spatiotemporal dynamics of microglia in the frontal cortex across the human lifespan (3^rd^ pcw – 75 years). We provide an unprecedented level of granularity and breadth, defining processes fundamentally different from the known development of murine microglia, which will influence how we study the role of these cells in humans. This was achieved by the collection of post-mortem specimens from a multitude of tissue resources, enabling us to conduct a cross-sectional study with an extensive scope. We define that microglia enter the telencephalon by the 4^th^ pcw, mirroring, with some delay, the colonisation of peripheral organs by other tissue-resident macrophages. After colonisation, microglia undergo very significant fluctuations in their cell density, characterised by newly identified wave-like patterns of proliferation followed by apoptosis. This is particularly evident at a key neurodevelopmental stage, the transition between embryonic and fetal life (9^th^ pcw). This pattern is corroborated by bulk RNAseq and single-cell RNAseq analyses and coincides with the appearance of the cortical plate and transient zones. Throughout embryonic, fetal and postnatal development, the population transits through different cycles of expansion and apoptosis-driven refinement, stabilising during infancy and being maintained by self-renewal across the adult and older ages.

The elucidation of microglial dynamics during human cortical development offers new avenues for examining how these cells contribute to neurodevelopmental disorders. These cells participate in brain wiring pre- and postnatally (Menassa and Gomez-Nicola, 2018; Paolicelli et al., 2011; Squarzoni et al., 2014) and are involved in the pathophysiology of autism spectrum disorder (Carroll et al., 2021; Tetreault et al., 2012), schizophrenia (Sekar et al., 2016), and intellectual disability (Coutinho et al., 2017). With our identification of critical temporal windows of the population’s expansion and refinement, we pave the way for novel uncharted territories in the field of neurodevelopmental disorders, which, with the current tissue resources in place, may begin to closely characterise in space and time how the population alters the neurodevelopmental environment. We know that targeting microglial dynamics may have a therapeutic potential when these cells go awry, and this is likely to extend to the treatment of neurodevelopmental disorders (Olmos-Alonso et al., 2016).

During our study we encountered several challenges, reflective of the scarcity of postmortem brain samples from key ages. For example, we could not obtain tissues from childhood and adolescence, a period of protracted intracortical myelination in the frontal and temporal cortices in humans (Deoni et al., 2015). As we know, microglia follow myelination very closely (Menassa and Gomez-Nicola, 2018; Verney et al., 2010); the dynamics of the population have yet to be carefully defined during the critical window of intracortical myelination in humans. Early adolescence is a dynamic period and is important for the onset of some neuropsychiatric disorders (Penzes et al., 2011). Additionally, as there are documented sex-specific differences in microglia (Lenz and McCarthy, 2015), hormonal changes during puberty are likely to influence microglial dynamics. Another challenge relates to our definition of migration, which did not allow us to distinguish between peripheral migration (into the brain) and within-brain migration. Microglia acquire a migratory phenotype subsequent to a proliferation cycle (Martín-Estebané et al., 2017) and our assessments here are of a migratory phenotype irrespective of contribution. We observed that within-brain migration varies according to the topography of transient layers defined by the type of neurogenetic process. Overall, migration contributes to the growth of the population at specific timepoints during development, thereafter, and once microglia are in the parenchyma and proliferating, they can acquire a migratory phenotype after each proliferative cycle to tile the anatomical local territory they are destined for.

Altogether, our findings identify key features of and relevant temporal windows for microglial population expansion and refinement across the human lifespan. This study provides a solid foundational map of the precise microglial spatiotemporal dynamics from early development through adulthood to healthy ageing. This resource will inform future research on how microglial cells participate in the neurodevelopmental landscape in humans and their relevance for neurodevelopmental disorders as they form part of the pathological signature of these conditions (Coutinho et al., 2017; Velmeshev et al., 2019).

## ACKNOWLEDGEMENTS

The research was funded by the Leverhulme Trust (RPG-2016-311), the Medical Research Council (MRC) (MR/P024572/1) and the Alfonso Martin-Escudero Foundation (fellowship to M.M-E).

We would like to thank the joint MRC/Wellcome Trust Human Developmental Biology Resource (HDBR) (grant # MR/006237/1) (Gerrelli et al., 2015; Lindsay et al., 2016), the Zagreb Research Brain Collection (Hrabac et al., 2018), and BRAIN UK, a collaboration between representatives from the University of Southampton, Plymouth Hospitals NHS Trust, the University of Bristol, and a network of many NHS neuropathology centres across the United Kingdom. BRAINUK is funded by the MRC and Brain Tumour Research. In this study, the following NHS neuropathology centres that are part of BRAIN UK participated: The South-Central Oxford NHS Trust (the Thomas Willis Oxford Brain Bank), the South-Central Hampshire NHS Trust (Southampton Brain Bank) and the Cambridgeshire NHS Trust (Cambridge Brain Bank). We also thank the East Scotland NHS Trust (Edinburgh Brain Bank) and the North Somerset and South Bristol NHS Trust (Southwest Dementia Brain Bank) for providing tissues for this study. We would like to thank the Research Cooperability Program of the Croatian Science Foundation funded by the European Union from the European Social Fund under the Operational Programme Efficient Human Resources 2014-2020 PSZ-2019-02-4710 (ZK). Many thanks to the Scientific Centre of Excellence for Basic, Clinical and Translational Neuroscience (project entitled Experimental and clinical research of hypoxic-ischemic damage in perinatal and adult brain; GA KK01.1.1.01.0007 funded by the European Union through the European regional development fund). We thank the Institutional Excellence in Higher Education Grant (FIKP) and Thematic Excellence Programme (National Research, Development and Innovation Office, Hungary), which supported the work of TT and IA. We would like to thank Steven Lisgo, Janet Kerwin, Tabitha Bloom, Carolyn Sloan, Chris-Anne McKenzie, Laura Palmer, Oliver Green, Poonam Singh, Danica Budinscak, Ana Bosak, Jenny Dewing, David Chatelet, Olaf Ansorge and Thomas Jacques for all their technical support in this project. We acknowledge the IRIDIS High Performance Computing Facility and associated support services at the University of Southampton towards the completion of this work.

## AUTHOR CONTRIBUTIONS

DAM and DG-N designed the study. DG-N secured the funding and supervised the project. DAM contacted the relevant brain banks, organised the collection, performed all histological experiments, scanned the slides, and analysed the data. MM-E performed the migration analysis. TT and IA constructed the heatmaps and provided expertise about case selection. ZK and IK provided neuroanatomical expertise and neurogenetic analysis of microglial findings. MC performed the bulk-RNA-seq analysis. TAOM performed the sc-RNA-seq analysis. JN assisted in the assessment of the neuropathology for case selection. LB-C and BT assisted with cell count analysis. DAM and DG-N wrote the manuscript. All authors contributed to drafting the manuscript.

## DECLARATION OF INTERESTS

The authors declare no conflict of interest.

## METHODS

### Human Tissues

Human developmental tissues were obtained through the joint Medical Research Council (MRC)/Wellcome Trust Human Developmental Biology Resource (HDBR) (Gerrelli et al., 2015), the Zagreb Research Brain Collection (Hrabac et al., 2018) and BRAINUK neuropathology centres (Fig 1; Supplementary table 1 for demographics of cases and acknowledgements for participating BRAINUK centres). Tissues were collected with appropriate maternal consent and approval for use in research. Ethical approval was obtained from the relevant ethics committees (Newcastle and North Tyneside NHS Health Authority Research Ethics Committee, Fulham NHS Health Authority Research Ethics Committee, South-Central Oxford C NHS Health Authority Research Ethics Committee and the South-Central Hampshire B NHS Health Authority Ethics Committee). Embryonic ages were estimated according to the Carnegie classification (CS) provided by the HDBR, CS23 being the last Carnegie stage equivalent to the 9^th^ pcw. Gestational age corresponds to the time elapsed between the first day of the last menstrual period and the day of delivery (Engle et al., 2004). We used here postconceptional age consistent with HDBR guidelines which is estimated at 2 weeks earlier than gestational age (Supplementary table 1). For fetal developmental stages (age>9 pcw until term), cases were aged by a neuropathologist according to clinical notes. When possible, developmental cases were sex-matched by timepoint. 63 developmental cases were initially sampled but only 52 developmental cases were included in the final study (n = 15 embryonic tissues; n = 37 fetal tissues; Fig 1 and supplementary table 1) between the late 3^rd^ pcw (CS10) and 38 pcw (term). We excluded cases due to poor immunoreactivity (poor antigenicity) or evidence of hypoxic injury identified in the clinical notes (Supplementary fig 1; supplementary table 1). Where possible, maternal data were provided (Supplementary table 2). Exclusion criteria, assessed against individual medical histories, included brain injury due to hypoxia-ischaemia or trauma, infection, or genetic mutations affecting brain structures. 71 postnatal cases were collected between 0 and 75 years of age with 45 of these cases in the final study. Early postnatal brain tissues (n = 24) aged between 0-2 years were obtained through BRAINUK. Adult brain tissues (n = 21) aged between 18-75 years were obtained with informed written consent and material approved for use for research purposes through the Zagreb Brain Collection, Edinburgh Brain Bank, the Cambridge Brain Bank and the Southwest Dementia Brain Bank. Ethical approvals were from the East Scotland NHS Research Ethics Service, the Cambridgeshire NHS Health Authority Research Ethics Committee and the North Somerset and South Bristol NHS Health Authority Research Ethics Committee, respectively. It was not possible to get sex-matching in postnatal cases (n = 45) and additional exclusion criteria included any neurological co-morbidity or a diagnosis of a connectivity disorder (e.g., autism spectrum disorder) (Supplementary table 3). We could not obtain samples from children aged between 3 and 17 years as these were scarce and, when present, the absence of consent prohibited the usage of these tissues for research.

All tissues were received as paraffin-embedded slices previously fixed in formalin phosphate buffer saline and sometimes with an added secondary fixative, methacarn (HDBR samples). Mid-sagittal embryonic sections of 8 μm thickness were obtained through the whole embryo (up to CS21) allowing the visualisation of the brain rudiment and organs such as the heart, liver and spleen. All other sections, from CS23 until the late postnatal ages were processed coronally through the frontal axis of the brain at a thickness of 8 μm.

### Anatomy and Neuropathology

Embryonic samples along the midsagittal axis included the telencephalic wall with its dorsal and ventral components. The frontal cortex (dorsolateral prefrontal, anterior cingulate) develops from the dorsal telencephalon (Bayer and Altman, 2006, 2008). After CS21, the medial ganglionic eminence (GE) in the ventral telencephalon becomes more prominent. From CS23 onwards, developmental sections were all along the coronal axis of the frontal lobe with a prominent medial GE, an expanding telencephalic wall featuring transitional zones (marginal zone (MZ), cortical plate (CP), pre-subplate (PSP), subplate (SP), intermediate zone (IZ), subventricular zone (SVZ), ventricular zone (VZ); supplementary fig 2). By 32 pcw, frontal cortex grey and the underlying white matters are developed, transitional zones have largely resolved, and brain structures begin to resemble those from adulthood. All sections were histochemically labelled with haematoxylin and eosin according to standard methods, assessed by a neuropathologist for any signs of tissue pathology prior to analysis such as local or generalised hypoxia, haemorrhage, gliosis or neuronal death. Developmental tissue sections were histochemically labelled with filtered cresyl violet for the Nissl substance to visualise transitional zones for anatomical delineation as specified elsewhere (Duque et al., 2016; Kostovic et al., 2002). The SP was visualised by histochemical labelling of the extracellular matrix using the Periodic Acid Schiff Reagent-Alcian Blue method as specified elsewhere (Kostovic, 2020; Kostovic et al., 2002). White matter was visualised using the Gallyas silver labelling method by means of physical development (Gallyas, 1979).

### Immunohistochemistry

For immunostaining, sections were placed in an oven at 60°C for 45 min, dewaxed in 100% xylene solution, rehydrated in decreasing graded absolute ethanol (in distilled water) solutions and washed in water and 0.1% TWEEN 20-phosphate buffer saline (PBS-T) solution. Heat-mediated antigen retrieval in 10 mM citrate buffer (pH=6.2) was subsequently performed on the sections in a microwave for 25 min. For early timepoints (CS10-CS13), 30 min antigen retrieval was performed using 0.5% sodium borohydride (213462, Sigma-Aldrich, UK). Then, sections were washed and/or cooled in cold tap water. Endogenous peroxidase and phosphatase activity in the tissues was quenched with a dual enzyme block from Dako (S2003, Agilent, UK). Sections were washed in 0.1% PBS-T and blocked with the relevant serum (goat, horse or rabbit) and 5% bovine serum albumin (BSA) in 0.2% PBS-T for one hour. Microglia were probed with a 24h incubation at 4°C with either a rabbit antihuman IBA1 antibody diluted in blocking solution (1:200, 019-19741, Wako chemicals, USA) or a goat anti-human IBA1 antibody (1:200, ab5076, Abcam, UK). We used additional microglial markers including mouse anti-MHC-II/HLA-DP/DQ/DR (1:250, M077501-2, Agilent, UK), rabbit anti-PU.1 (1:200, 2258S, Cell Signalling, UK) and biotinylated RCA1 (1:1000, B10855, Vector Labs, UK). Mouse anti-Ki67 (1:400, M724029-2, Agilent, UK) was used to label proliferative cells and rabbit anti-cleaved-Caspase3 (1:40, 9664S, Cell Signalling, UK) was used to label apoptotic cells. Following incubation with the primary antibody, sections were washed with 0.1% PBS-T and incubated with the appropriate secondary antibodies. Secondary antibody double labelling was through an ImmPress Duet double staining anti-mouse HRP (brown)/anti-rabbit AP (magenta) kit (MP7724, Vector Laboratories, UK) according to the manufacturer’s instructions; horse anti-rabbit HRP using DAB chromogen with Nickel (SK100, Vector Laboratories, UK); horse anti-rabbit AP (blue) using a BCIP/NBT substrate kit (SK5400, Vector Laboratories, UK); rabbit anti-mouse secondary using the ImmPACT DAB-EqV HRP (brown) substrate kit (SK4103, Vector Laboratories, UK); signal amplification after species-specific secondary biotinylated antibody incubation of two hours was done using an avidin/biotin-based peroxidase system (PK6100, ABC Vectastain Elite kit, Vector Laboratories, UK). Chromogen development was sequential with double labelling. For triple labelling in brightfield, an additional DAB chromogen development was used, sections were re-quenched, re-blocked and probed with the relevant primary and secondary antibodies (bound to HRP) then developed with DAB + Nickel (SK4100, Vector Laboratories, UK). Chromogen development reactions were halted with distilled water, sections were washed with 0.1%PBS-T for 15 min then counterstained with methyl green (H3042, Vector Laboratories, UK) for 10 min or haematoxylin QS (H3404-100, Vector Laboratories, UK) for 30 seconds diluted 1:3 in distilled water. Sections were subsequently washed in distilled water for 5 min, then dehydrated in increasing gradients of absolute ethanol, cleared in xylene for 15 min, coverslipped with permanent mounting medium (DPX), dried for 24 hours then cleaned for scanning.

### Image analysis

Slides were digitised using a VS110 high-throughput virtual microscopy system (Olympus, Japan), an Aperio Scanscope AT Turbo system (Leica Biosystems, UK) or a 2.0 RS Nanozoomer high-throughput system (Hamamatsu Photonics, Japan). Pixel resolution was 172.35 nm/pixel in x, y and z. From the scans, regions of interest were extracted, and brain/layer thicknesses were measured in the relevant slide viewers (VS110-Desktop (v4), Aperio ImageScope (v12.4.3), NDP.view2). Images were processed using Fiji (Schindelin et al., 2012). The analysis pipeline included re-scaling the images, deconvoluting the signal and calculating a cell density (in cells/mm^2^) based on absolute counts until CS23. Beyond that age, between 500-1000 cells/case were sampled as deemed sufficient to provide a representative value of brain IBA1 densities consistent with unbiased sampling methods (Herculano-Houzel et al., 2015). The proliferative index -I- of microglia was calculated as follows: I = (number of double positive (IBA1/Ki67) cells/total number of IBA1 cells) x 100 (see also (Askew et al., 2017)). Double positives were confirmed by deconvolution (Supplementary fig 3). Prenatally, we calculated fold change values between subsequent ages based on overall cortical wall or individual layer thickness measurements to account for brain/layer growth rates during development. Postnatally, we collected brain weight data (when available) from the clinical notes of analysed cases and calculated the fold change increase in brain weight between subsequent ages and applied this correction to our density data. Proliferation index was independent of the changes in brain growth rates. Individual layers were manually traced in Fiji using a macro (Supplementary data 1) and a cell density was calculated as N_D_ = (Number of cells/area (mm^2^)).

### Migration analysis

The migratory phenotype of microglia (IBA1^+^ cells) was assessed using morphometric analysis in Fiji (Schindelin et al., 2012). The circularity parameter was used to select out the migratory phenotype in each layer: if circularity<0.3, a cell would be considered migratory and if circularity>0.3, it would be non-migratory (Supplementary fig 4A-C). We also assessed the type of migration (radial or tangential) by measuring the angle formed between the major axis of the cell and the corresponding layer plane (Supplementary fig 4C-D), considering an angle<45 degrees as tangential migration and an angle>45 degrees as radial migration. Developmental timepoints considered were selected based on the cell density profile whereby an increase could not be explained by proliferation alone. We studied migration between 5 and 26 pcw (n=2/timepoint).

### Histological heatmaps

For the visualization of cell densities as heatmaps, cortical columns per stage were manually annotated in Aperio ImageScope software ((v12.4.3), LeicaBiosystems, IL, USA) (Supplementary figure 5). At least 2 cases by timepoint were analysed and the most representative column was used for visualisation in the final spatiotemporal profile. IBA1 and IBA1/Ki67 double-positive cells were analysed separately. Coordinates of the annotated cells were exported into Quantum Geographic Information System QGIS (v2.18.3, Hannover, Germany), a spatial analysis software. The heatmap module in QGIS with metric projection EOV23700 was used. We used a sampling radius of 150 μm, with a maximum value of 4, which meant that on the red area of a computed heatmap in a (0,15 mm^2^)π area of a circle, at least 4 cells could be located. Subsequently, proportionally extrapolated values were used to estimate the density in cells/mm^2^. Heatmaps were exported and superimposed onto their respective immunohistochemically labelled spatial maps using Gimp 2.10.22 annotating the anatomical boundaries.

### Bulk RNA-sequencing analysis

We sourced published lists of genes from adult and adolescent human cortical microglia (Galatro et al., 2017; Gosselin et al., 2017). 906 microglia genes were identified (Supplementary data 2) and were mapped onto an available bulk-RNAseq dataset from the HDBR. We obtained access to n = 251 samples of tissues ranging in age from 7 to 17 pcw. Four anatomical regions were considered: telencephalon (n = 94), cerebellum (n = 79), choroid plexus (n = 28) and midbrain (n = 50). Full details of the source, collection, preparation and sequencing of human fetal RNA samples have been described previously (Gerrelli et al., 2015; Lindsay et al., 2016) (https://www.hdbr.org/expression/). Data were downloaded from SRA as FASTQ files. Quality control was carried out using Trimmomatic to remove poor quality bases, reads with too many poor-quality bases and short reads using the following *settings: ILLUMINACLIP:/local/software/trimmomatic/0.32/adapters/TruSeq3-PE-2.fa:2:30:10 LEADING:5 TRAILING:5 SLIDINGWINDOW:4:15 MINLEN:72.* After this, unpaired reads were excluded from further analysis. STAR v2.5.2b (Dobin et al., 2013) was used to map reads back to the GRCh38 genome using settings --outFilterMismatchNmax 10 --outFilterMismatchNoverReadLmax 0.05 and the result outputted as .sam files, which were sorted using Samtools (v1.1, (Li et al., 2009)). Read counts were calculated per gene using HTSeq count (in the HTSeq v0.6.1 package; (Anders et al., 2015)) and the GRCh38 general feature format file. Differential gene expression (DEGs) analysis was carried out using EdgeR (Robinson et al., 2010) in Trinity (v2.4.0) (Haas et al., 2013) on raw read counts. Heatmaps were generated using *analyze_diff_expr.pl* in Trinity (Haas et al., 2013) and the TMM-normalised counts. Gene Ontology (GO) terms over-represented in the list of DE genes were identified using the plug-in available at the Gene Ontology resource.

### Single-cell RNA-sequencing analysis

To validate our histological and bulk-RNAseq findings, we analysed 4 datasets from recently published developmental single-cell RNA-sequencing studies of human brain cells (Bian et al., 2020; Cao et al., 2020; Fan et al., 2020; Kracht et al., 2020) and generated an integrated dataset spanning 3 to 24 pcw (Bian et al., 2020; Kracht et al., 2020). We accessed the genecell count matrix and cell annotation matrix data and used Seurat (v3.2.2) for all analyses (Stuart et al., 2019). Guided by the original authors’ annotations, we enriched for microglial cells by selecting for “Immune”, “Mac_1”, “Mac_2”, “Mac_3”, “Mac_4”, and “Microglia”-annotated clusters. We then utilized the 3 x Mean Absolute Deviation (MAD) for outlier cutoff across 4 parameters where available: nCount_RNA, nFeature_RNA, percent.mt, percent.rb (Daniszewski et al., 2018; Kracht et al., 2020; Waise et al., 2019). After quality control, the final integrated dataset used for our analyses contained 24,751 transcriptomes, 8,117 of which were nuclei. Standard Seurat SCTransform integration was performed, selecting 3,000 features for anchor identification and integration and regressing for nCount_RNA, nFeature_RNA, percent.mt, and/or percent.rb (Supplementary fig 6). We detected immediate-early gene expression suggestive of dissociation-induced artefacts. However, a discussion of these effects in human tissue was beyond the scope of the study and were not selected for regression. 10 principal components were selected for dimensionality reduction and combined with a resolution of 0.5, to visualize transcriptional heterogeneity across human development spanning 3 to 24 pcw. Cell cycle phase was determined with ‘CellCycleScoring’ to identify actively proliferating cells. Only ages with more than 50 cells were selected for the proliferation wave signature. Data were visualized using DimPlot, FeaturePlot, VlnPlot and DoHeatmap functions, and differential expression analysis was performed with MAST (Finak et al., 2015).

### Statistical analysis

Statistical analysis and visualisations were done in RStudio and GraphPad Prism. We have predominantly used non-parametric techniques in our analyses and therefore, made no assumptions about data distributions or homoscedasticity. Non-parametric correlations using Spearman’s tested associations between brain weight, layer thicknesses and age. Sex differences were assessed using cumulative distribution plots with the 2-sample Kolmogorov-Smirnov test. Friedman’s test was used to test for differences between layers within matched data. Wilcoxon test was used to test differences between matched data (liver and brain from the same cases). We also fitted non-parametric regression lines to density and proliferation data using the following parameters: a Loess regression function with a medium number of 10 points followed by a smoothing spline function with 6-8 knots as smoothing factors, which were both recommended by Graphpad Prism. We tested centred polynomial models of up to the maximum recommended order (6^th^ order) to model our data but these could explain at most 60% of the variance. Therefore, these were not suitable and we opted for presenting all datapoints instead and applying non-parametric functions that follow the data trend with no assumptions about a model that could fit all data. Increasing the order of polynomial models may have improved variance but would have resulted in overfitting. Furthermore, biological fluctuations that we report in the microglial population underlie the difficulty of finding one model that fits all data. To test for statistical significance between temporal windows for density and proliferation in the cortex prenatally, we compared mean values between temporal windows at the peaks and troughs of the fitted regression lines using a non-parametric Kruskal-Wallis test corrected for multiple comparisons using Graphpad’s recommended Dunn’s test for multiple comparisons between mean-ranks and report throughout the adjusted corrected p-values. For infancy and adult stages, we used the Mann-Whitney U non-parametric test to report significant differences between means. We acknowledge that we were limited by the number of samples in late infancy and therefore, would require a larger dataset to better define the selection phase. However, collectively, proliferation and density values clearly show that the period of birth to late infancy is significantly different to the adult stage. For RNA-seq data (single cell and bulk), p-values, including those related to analysis of DEGs, were corrected for FDR in EdgeR using the Benjamini-Hochberg correction. Volcano plots and heatmaps were plotted in RStudio. Hierarchical clustering was carried out in Trinity (Haas et al., 2013). Histological heatmaps were plotted using QGIS spatial analysis software as elaborated upon in the relevant methods section.

## Notes

### Competing Interest Statement

The authors have declared no competing interest.

