## Supplemental Information for "Spatiotemporal dynamics of human microglia are linked with brain developmental processes across the lifespan"

**Index:**

Figs. S1 to S10

Additional supplementary files titles

**SUPPLEMENTARY FIGURES**

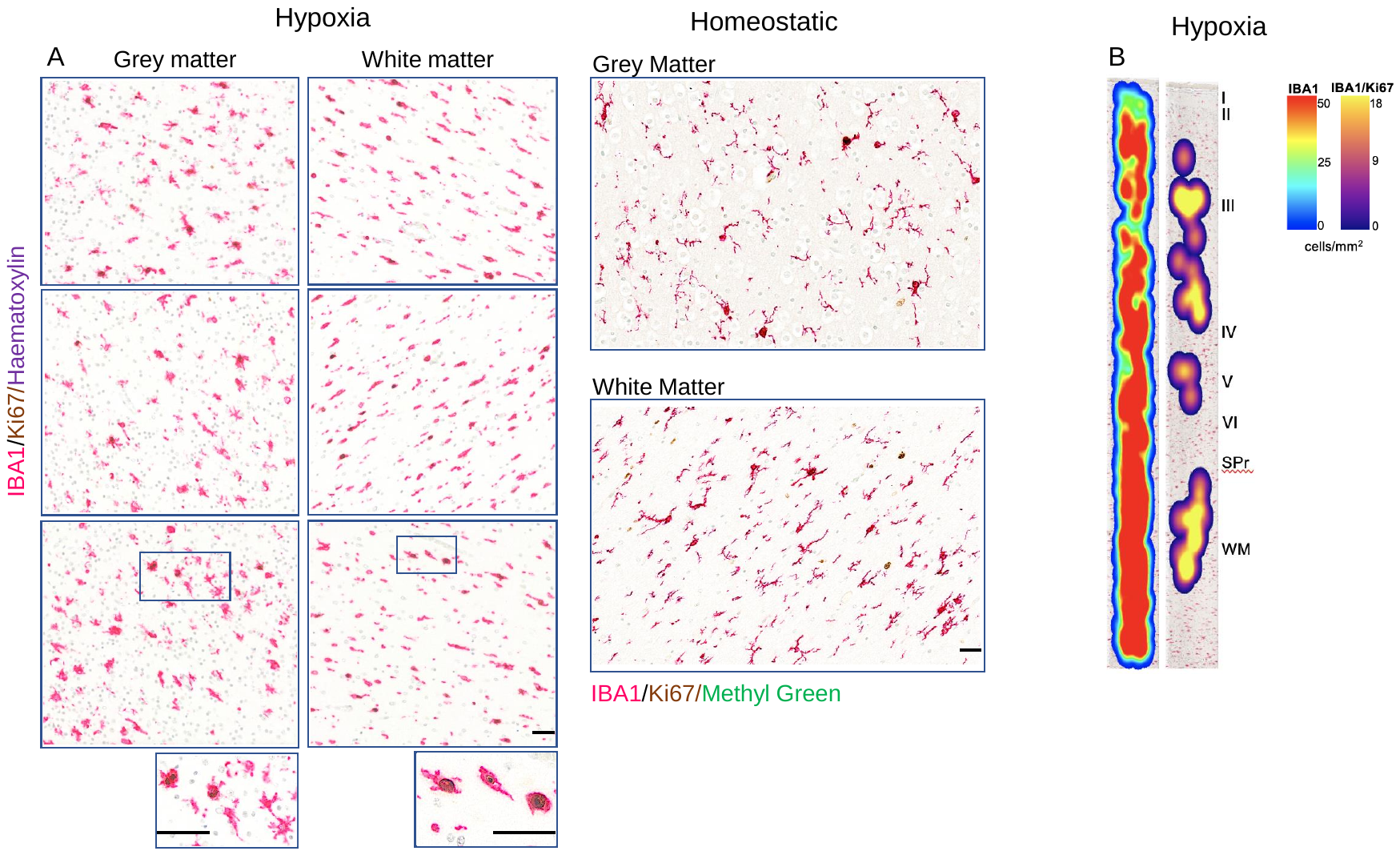

**Supplementary figure 1.** **Reaction of microglia to hypoxia.**

(A) Representative photomicrographs of microglia reactive to hypoxic-ischaemic injury in a 38 postconceptional term grey and white matters (Fetus 52 in table 1) and homeostatic microglial ramified morphology with a lower proliferation rate in the absence of hypoxic injury in a 38 postconceptional week case included in our study Note the amoeboid shape and higher proliferation rate compared to homeostatic microglia. (B) Representative heatmap of proliferation and density along the cortical column. *SPr*: subplate remnant; *WM:* white matter. Scale bar: 50μm.

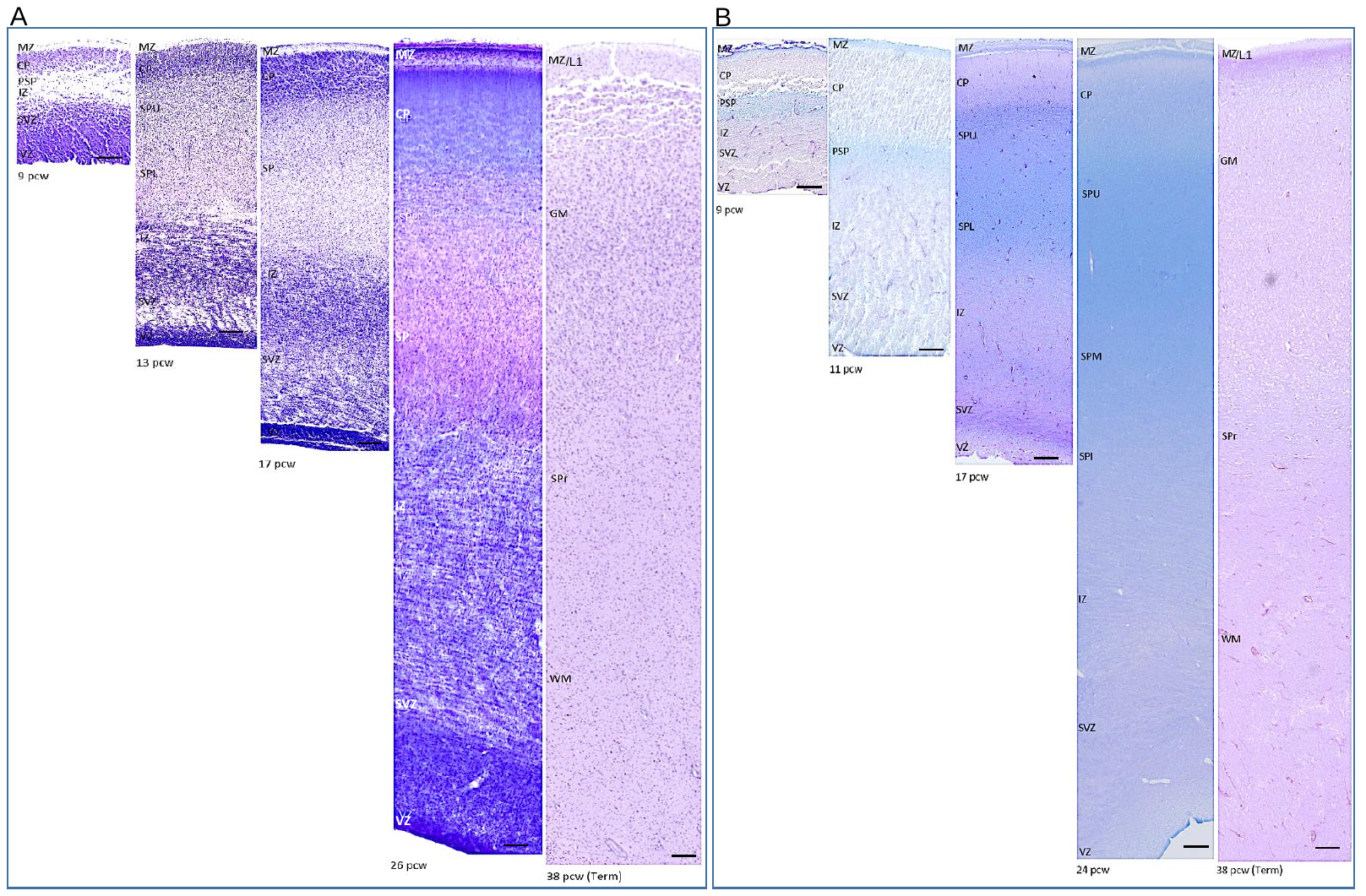

**Supplementary figure 2.** **Delineation of anatomical boundaries in the developing cortex between 9 and 38 postconceptional weeks.**

(A) Nissl histochemistry of transient cortical zones. (B) Periodic-Acid Schiff/Alcian Blue histochemistry for marking the subplate in transient cortical zones. *CP*: cortical plate; *GM:* Grey matter; *IZ*: intermediate zone; *MZ:* marginal zone; *PSP*: presubplate; *pcw*: postconceptional week; *SP*: subplate; *SPU:* subplate upper portion; *SPM*: subplate middle portion; *SPI:* subplate inferior portion; *SPL*: subplate lower portion; *SPr*: subplate remnant; *SVZ*: subventricular zone; *VZ*: ventricular zone; *WM:* white matter. Scale bar: 100μm.

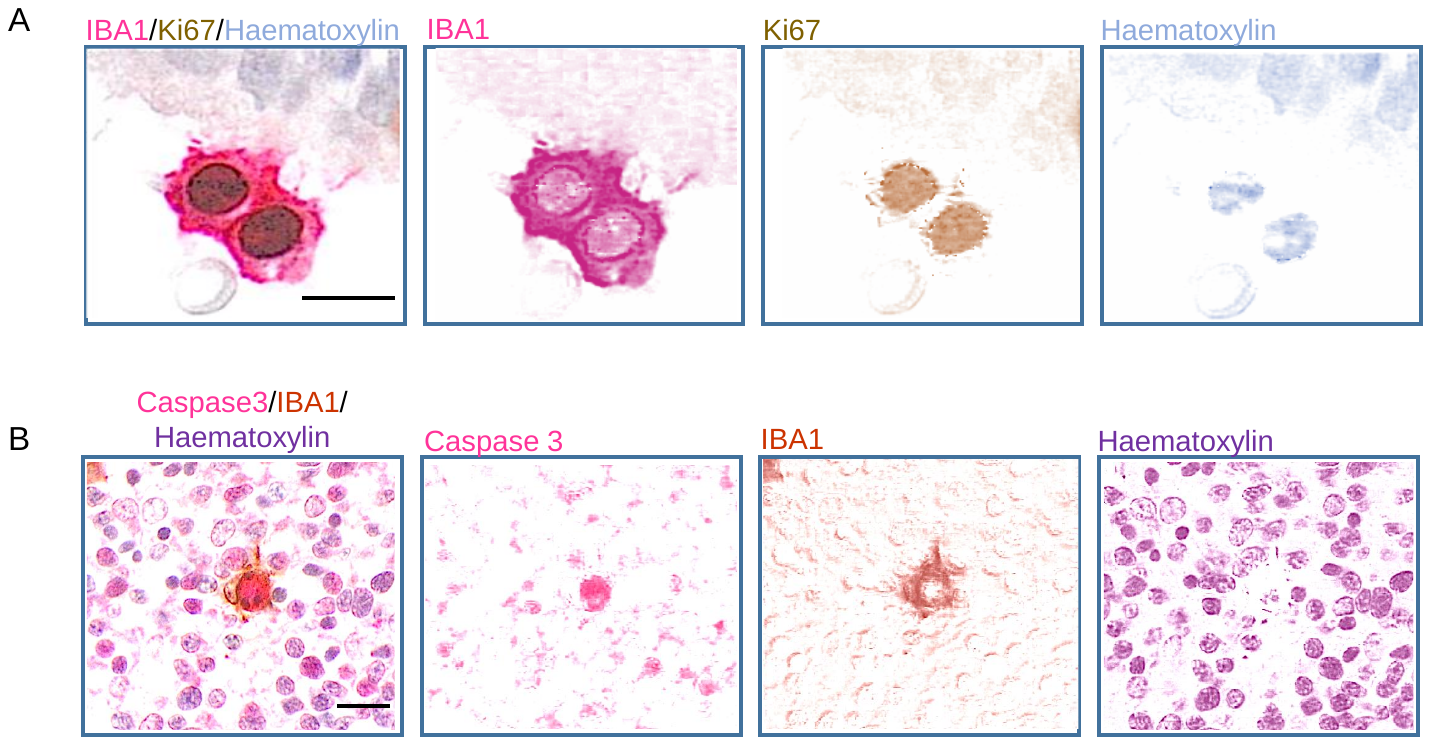

**Supplementary figure 3.** **Deconvolution of signal in brightfield.**

(A) Example of colour deconvolution in brightfield for proliferative microglia (IBA1/Ki67/Haematoxylin). (B) Example of colour deconvolution in brightfield for cell death (Caspase 3/IBA1/Haematoxylin). Scale bar: 10μm.

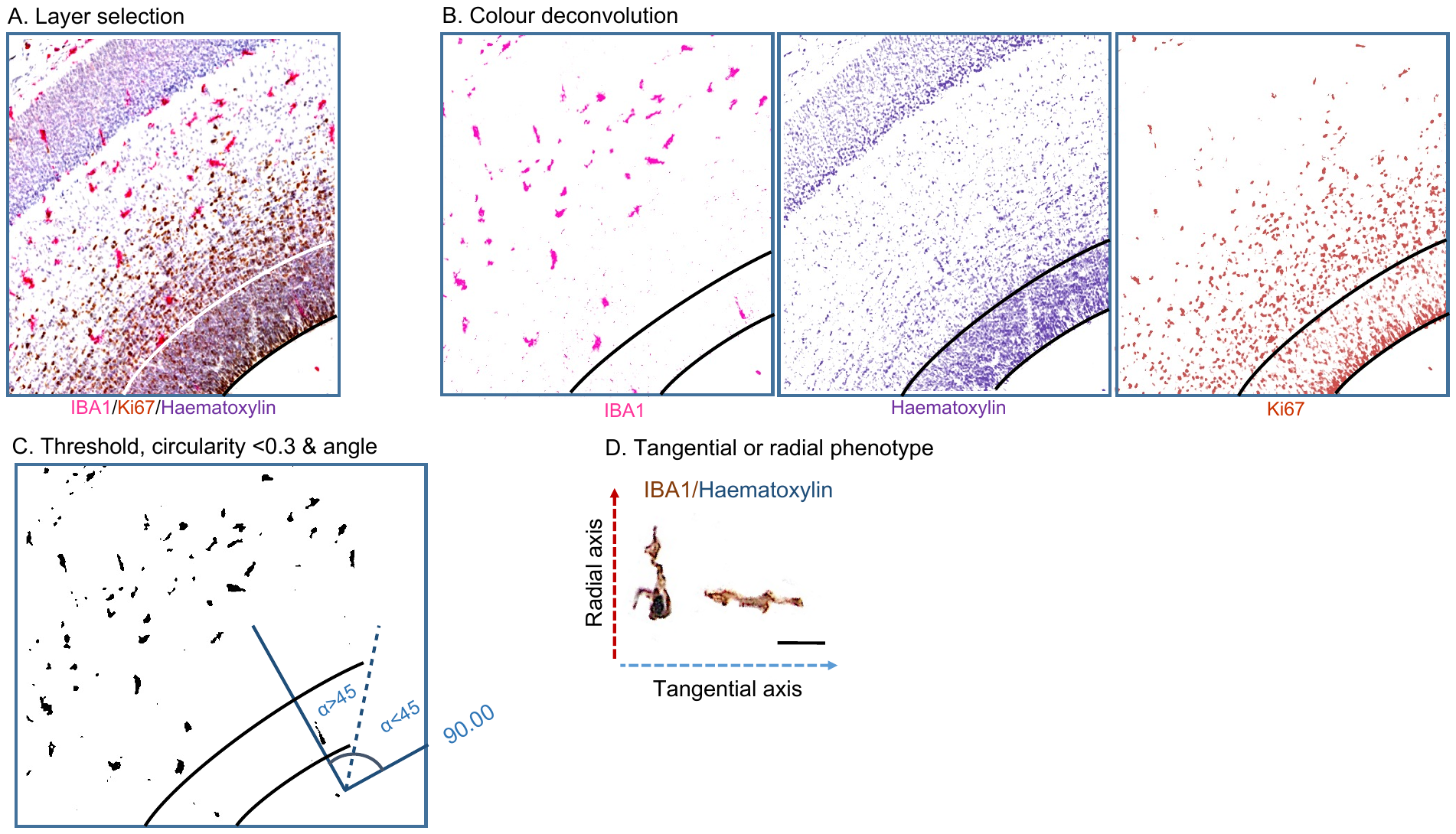

**Supplementary figure 4.** **Migration analysis workflow.**

(A) Layer selection. (B) Deconvolution. (C) Thresholded photomicrograph with particle selection according to a circularity criterion <0.3 for a migratory cell phenotype. The angle is recorded between the major axis of the cell and the layer plane with α<45 for tangential and α>45 for radial migrations. (D) Tangential and radial microglial phenotypes. Scale bar: 30μm.

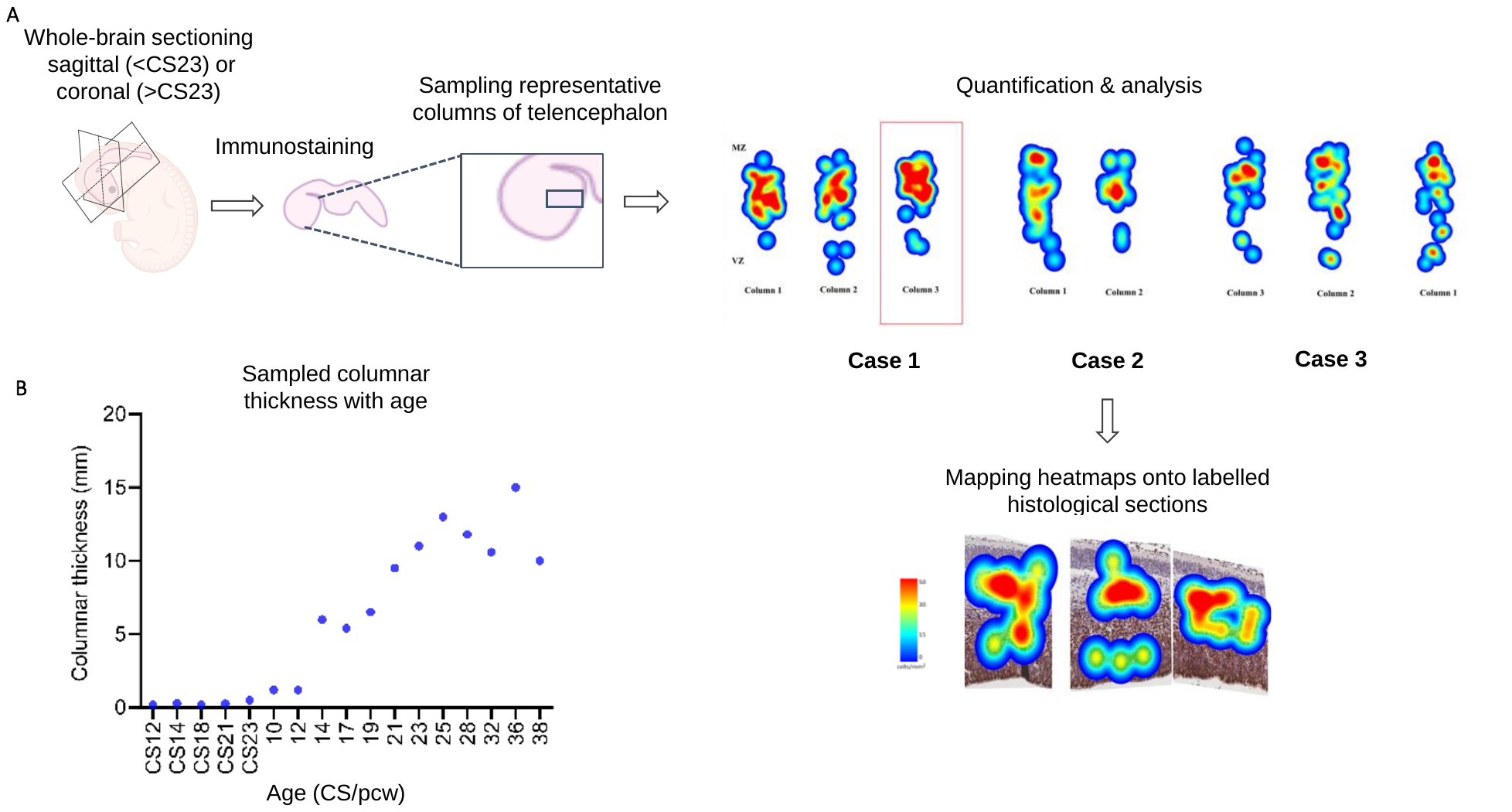

**Supplementary figure 5.** **Heatmap analysis.**

(A) Workflow of image processing to obtain heatmaps from cortical columns. (B) Columnar thickness measurements for heatmaps against age. *CS:* Carnegie stages.

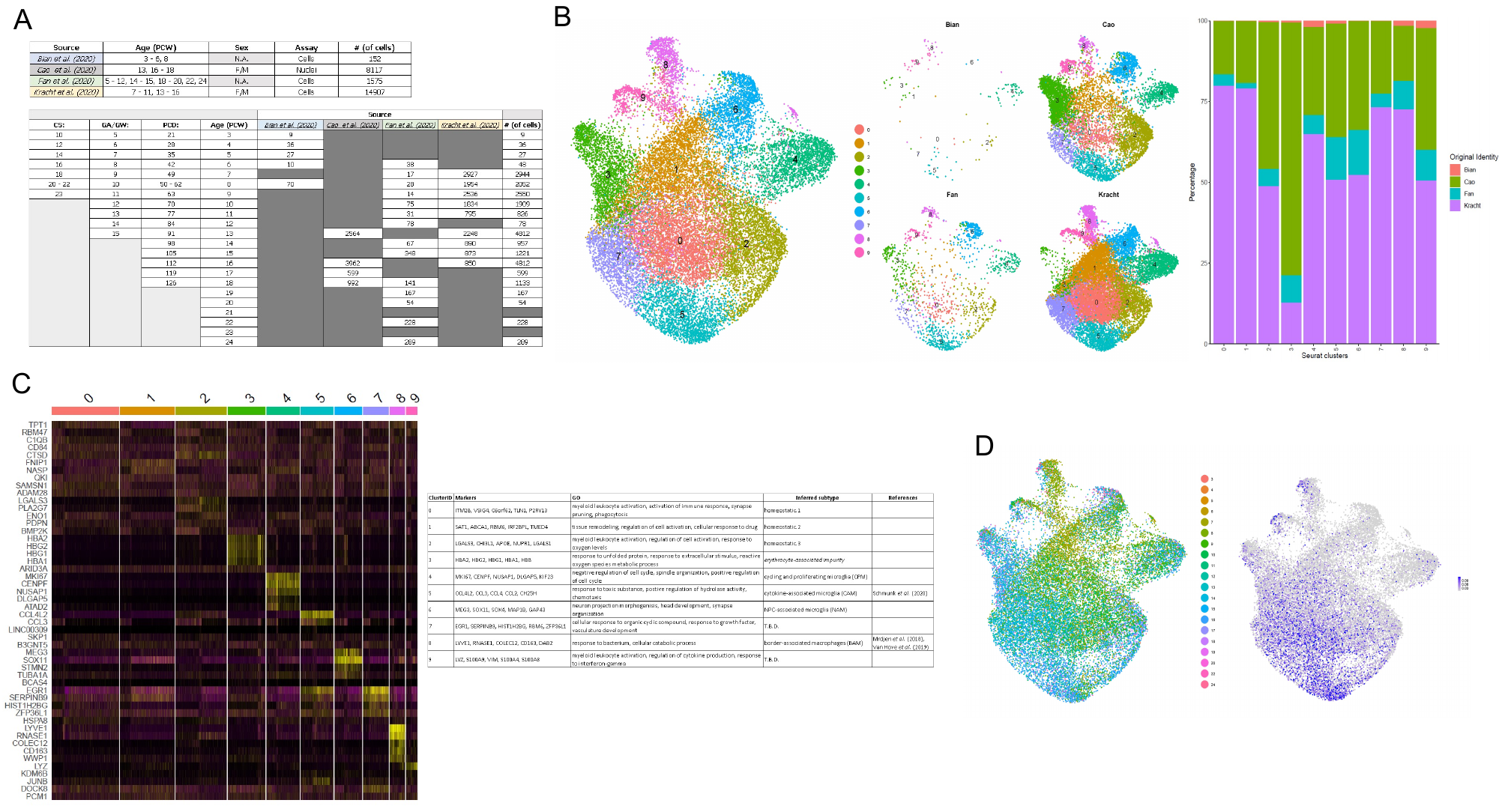

**Supplementary figure 6.** **Single-cell RNA-seq analysis.**

(A) Summary table of scRNAseq datasets (i.e. sources) (*22, 23, 43, 44*), spanning 3-24 postconceptional weeks, used in this study, their respective age-ranges, sex, the single-cell assay and the number of cells. (B) Transcriptional signatures and relative contributions across clusters. Cell contributions of each dataset to all clusters illustrates effective anchoring between datasets, albeit their underlying proportions differ. e.g. cluster 3 is enriched for cells that derive from [43], suggestive of a nuclei-specific signature less common in cell-based assays. (C) Transcriptionally distinct clusters were identified in our integrated object. The heatmap displays the top 5 DEG of each cluster, as identified with the FindAllMarkers function (MAST). Several developmental homeostatic microglia were identified (clusters 0 – 2), as well as several previously identified microglial subtypes (CPM, CAM, BAM); cluster 4, 5 and 8, respectively. Of note, two clusters displayed markers not commonly associated with microglia (i.e. cluster 3 and 6). Cluster 3 featured erythrocyte-associated genes, and was enriched in [43], suggestive of impurities in cell preparation more commonly linked to nuclei assays. Cluster 6 is associated with NPC-linked genes. Like cluster 3, cells in cluster 6 are likely to indicate impurities in preparation; or capture a novel microglial subtype that restricts the progenitor pool. (D) Microglial heterogeneity corresponds to adolescence and aged microglial signatures [35,36]. A plot by age (left) shows an age-dependent effect on heterogeneity. From the 906 genes identified in our bulk RNAseq dataset, 297 were detected in the scRNAseq dataset and illustrated by a FeaturePlot (right). A relative enrichment of the microglial signature was noted in late developmental stages, supportive of microglial maturation during gestational development   *CS*: Carnegie stage; *GW/GA*: gestational week/age; *PCD:* postconceptional day; *PCW*: postconceptional week.

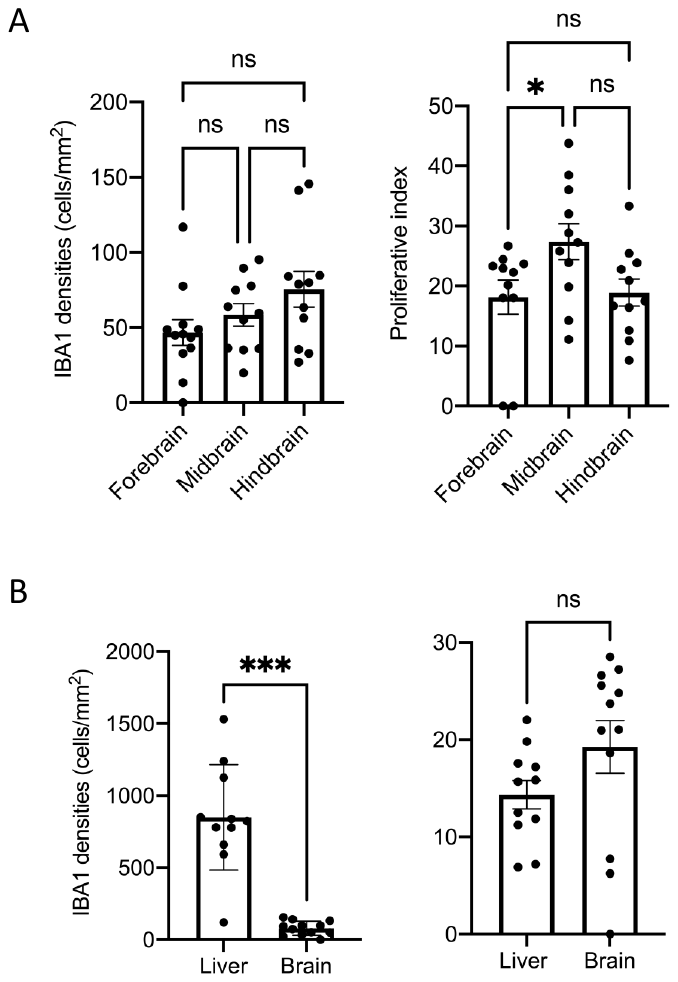

**Supplementary figure 7.** **Embryonic regional proliferation and densities.**

(A) IBA1 densities and proliferation between brain regions (Friedman’s test corrected for multiple comparisons using Dunn’s test; p<0.01) and (B) between the brain (n=15) and the liver (n=11) (Wilcoxon-matched-pairs, p<0.001).

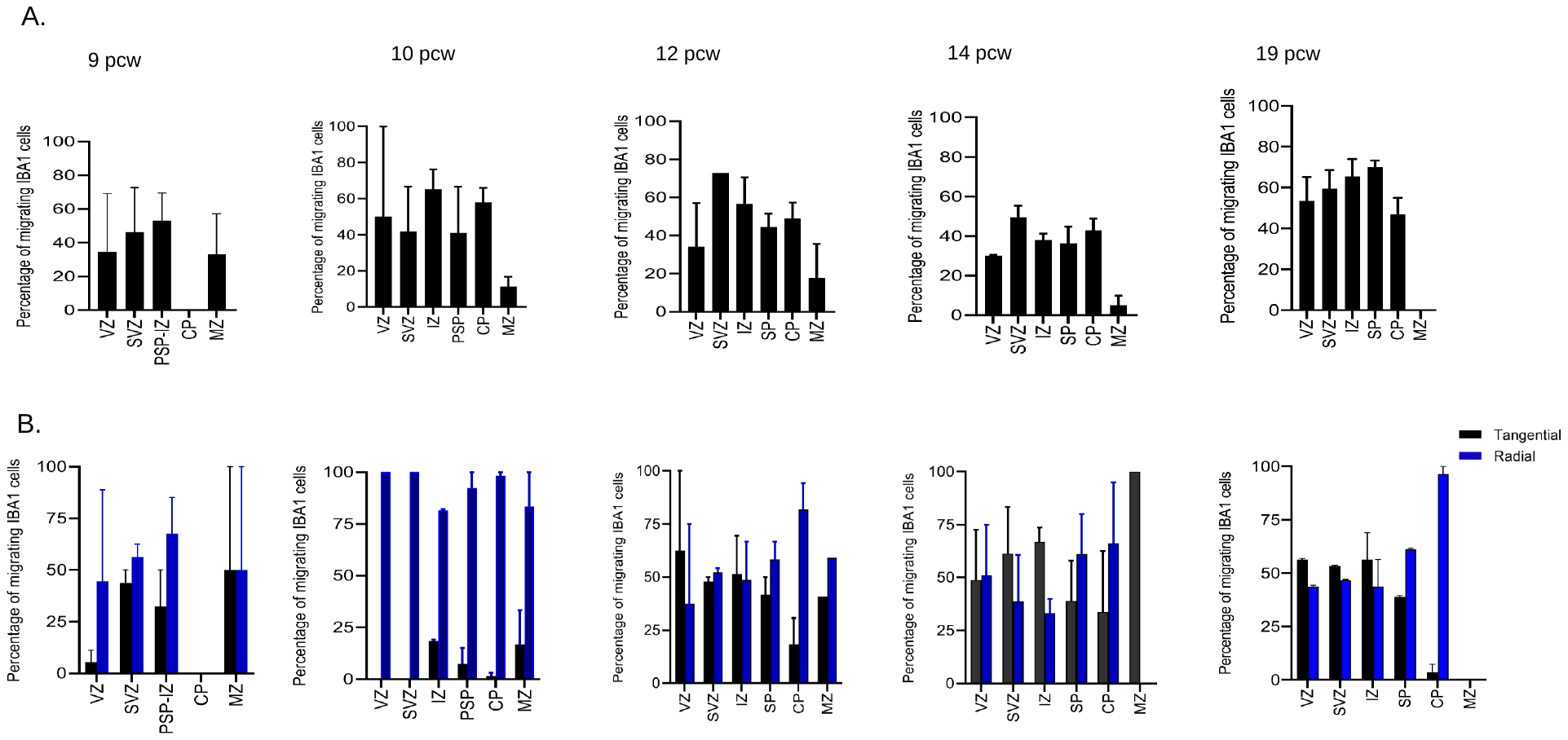

**Supplementary figure 8. Migratory microglial profile and phenotype during development.**

(A) Percentage of IBA1 cells migrating into the various transient zones (n=2 for each timepoint). At 9 pcw, the CP has no migrating microglia whilst most layers have substantial numbers of migrating cells. From 10-14 pcw, migrating cells are found in all layers and by 19 pcw, migrating cells can no longer be seen present in the MZ. (B) Type of migration in each transient layer (n=2 for each timepoint). Migration is radial and/or tangential depending on each layer across 9-19 pcw. *CP :* cortical plate ; *IZ :* intermediate zone ; *MZ :* marginal zone ; *PSP* : presubplate; *pcw*: postconceptional week; *SP*: subplate;; *SVZ*: subventricular zone; *VZ*: ventricular zone.

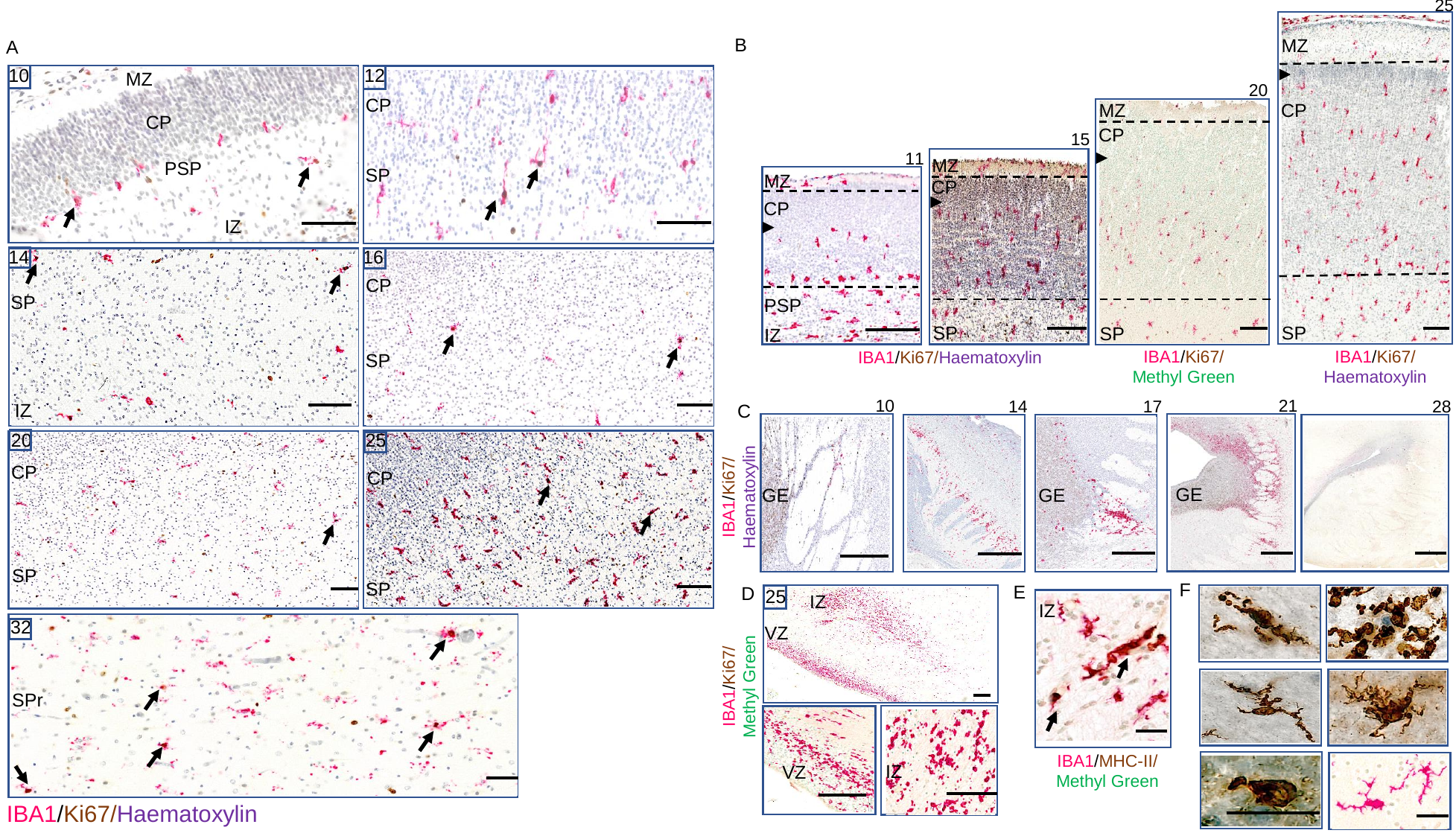

**Supplementary figure 9.** **Descriptive account of regional findings across development.**

(A) Proliferation of microglial cells is highest in the subplate region across gestation. Scale bar: 100 μm (B) Microglial cells invade the upper portion of the cortical plate by 25 pcw and spare it before that time. Scale bar: 100 μm (C) Ventral telencephalon microglia cluster around the internal and external capsules. These clusters increase during gestation and disappear by 28 pcw. Scale bar: 500 μm (D) Microglial hotspots can be seen in the intermediate zone during most of gestation and resolve by the 32^nd^ pcw. Scale bar: 100 μm (E) Microglia are positive for MHCII. Scale bar: 75 μm (E) and towards the late third trimester, cluster in the ventricular zone. (F) Range of microglial morphologies from amoeboid to intermediate and ramified mature types can be seen in transient layers developmentally. Scale bar: 75 μm. *CP :* cortical plate; *GE:* ganglionic eminence; *IZ :* intermediate zone ; *MZ :* marginal zone ; *PSP* : presubplate; *pcw*: postconceptional week; *SP*: subplate;; *SVZ*: subventricular zone; *VZ*: ventricular zone.

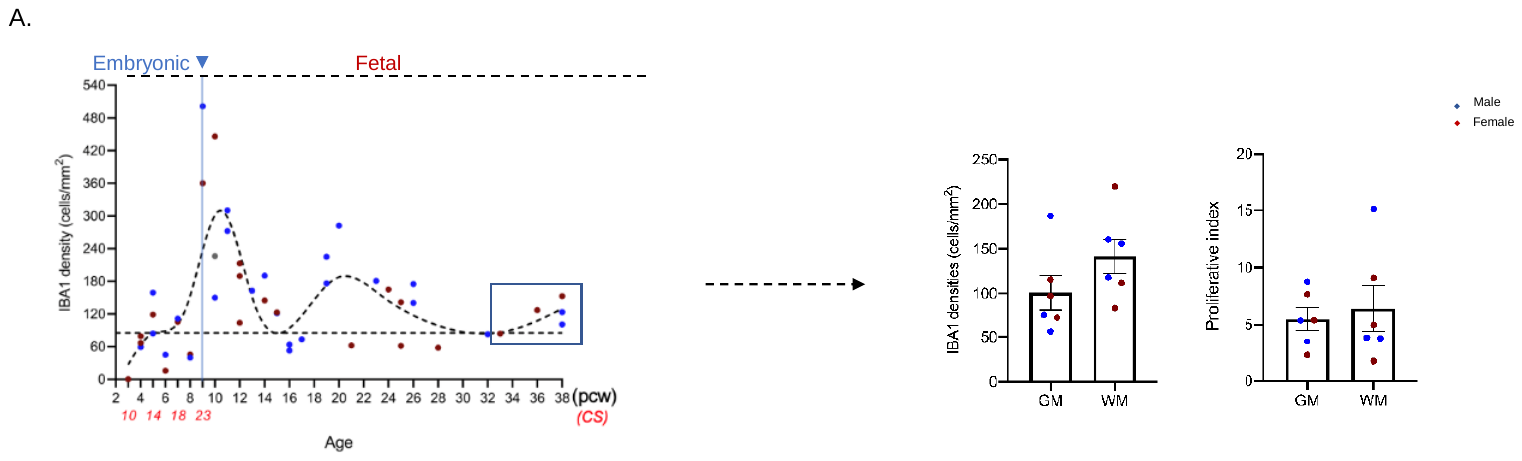

**Supplementary figure 10.** **Regional microglial differences during development.** (A) Grey and white matter densities and proliferation from 32 postconceptional weeks to birth. The neocortex has 6-layers by then and there is a clear distinction between grey and white matters similarly to the postnatal age.

**Supplementary table 1.** **Demographics of developmental cases.**

**Supplementary table 2.** **Maternal demographics.**

**Supplementary table 3.** **Demographics of postnatal cases.**

**Supplementary data 1.** **Code for region of interest histological analysis.**

**Supplementary data 2.** **RNAseq gene lists.**
