## Supplemental File 1 for "Spatiotemporal dynamics of human microglia are linked with brain developmental processes across the lifespan"

```
dir1 = getDirectory("choose the folder with the images");
dir2 = getDirectory("Choose a folder for the results");
list = getFileList(dir1);
Array.sort(list);

for(i=0; i<list.length; i++){
	open(dir1+list[i]);
	titleImage = getTitle();
	nameImage = File.nameWithoutExtension;
	run("Properties...", "channels=1 slices=1 frames=1 
unit=micron pixel_width=0.17235 pixel_height=0.17235 
voxel_depth=0.17235");
	getPixelSize(unit, pixelWidth, pixelHeight);
	layerNb = getNumber("how many tissue layers in this 
image:", 1);
	run("Duplicate...", "title=Duplicate");
	titleDupli = getTitle();
	for(j=1; j<=layerNb; j++){
		setTool("polygon");
		waitForUser("Layers"+j, "Trace the limits of the 
layer, then press OK");
		roiManager("Add");
		setForegroundColor(0, 0, 0);
		run("Fill", "slice");
	}
	selectWindow(titleDupli);
	close();
	selectWindow(titleImage);
	run("Duplicate...", "title=Duplicate");
	run("Properties...", "channels=1 slices=1 frames=1 
unit="+unit+" pixel_width="+pixelWidth+" 
pixel_height="+pixelHeight+" voxel_depth=1");
	run("From ROI Manager");
	run("Labels...", "color=white font=12 show draw");
	saveAs("TIFF", dir2+nameImage+"_layerOverlay.tif");
	close();
	roiManager("Save", dir2+nameImage+"_ROIset.zip");
	selectWindow(titleImage);
	run("Colour Deconvolution", "vectors=[FastRed FastBlue 
DAB]");
	selectWindow(titleImage+"-(Colour_3)");
	close();
	selectWindow(titleImage+"-(Colour_2)");
	close();
	selectWindow("Colour Deconvolution");
	close();
	selectWindow(titleImage+"-(Colour_1)");
	run("Properties...", "channels=1 slices=1 frames=1 
unit="+unit+" pixel_width="+pixelWidth+" 
pixel_height="+pixelHeight+" voxel_depth=1");
	run("Set Measurements...", "area area_fraction limit 
display redirect=None decimal=3");
	roiNb = roiManager("count");
	for(n=0; n<roiNb; n++){
```

```
                roiManager("Select", n);
                run("Duplicate...", "title="+nameImage+"_ROI"+n+1);
                setAutoThreshold("Default");
                //run("Threshold...");
                setThreshold(0, 255); // threshold the whole image
                run("Measure"); // measure the area of thresholded
tissue within the drawn ROI
                setThreshold(0, 170); // threshold the cells in the
image
                run("Measure"); // measure the areas of the
thresholded cells within the ROI
                setOption("BlackBackground", false);
                run("Convert to Mask");
                roiManager("Select", n);
                run("Erode");
                run("Dilate");
                run("Analyze Particles...", "summarize");
                saveAs("Jpeg", dir2+nameImage+"_Layer"+n+1+".jpg");
                close();
        }
        selectWindow(titleImage+"-(Colour_1)");
        close();
        selectWindow(titleImage);
        close();
        roiManager("reset");
}
selectWindow("ROI Manager");
run("Close");
selectWindow("Results");
saveAs("Results", dir2+nameImage+"_Results.csv");
run("Close");
selectWindow("Summary");
saveAs("Results", dir2+nameImage+"_Results_count.csv");
run("Close");
```
