## Supplemental Table 1 for "Spatiotemporal dynamics of human microglia are linked with brain developmental processes across the lifespan"

| **Supplementary Table 1: Developmental demographics** | | | | | | | | | | | | | |
| --- | --- | --- | --- | --- | --- | --- | --- | --- | --- | --- | --- | --- | --- |
| **Case** | **GA (CS)** | **PCW** | **Sex** | | **PMI (h)** | | | | | **BW (g)** | **Cause of Death** | | **Histology** |
| Embryo | 5 (10) | Late 3^rd^ | F | | 3.67 | | | | | n/a | Elective termination | | Normal |
| Embryo | 6 (12) | 4 | M | | 3 | | | | | n/a | Elective termination | | Normal |
| Embryo | 6 (12) | 4 | F | | 3.92 | | | | | n/a | Elective termination | | Normal |
| Embryo | 6 (12) | 4 | F | | 4.67 | | | | | n/a | Elective termination | | Normal |
| Embryo | 7 (14) | 5 | M | | 7 | | | | | n/a | Elective termination | | Normal |
| Embryo | 7 (14) | 5 | F | | 6.42 | | | | | n/a | Elective termination | | Normal |
| Embryo | 7 (14) | 5 | M | | n/k | | | | | n/a | Elective termination | | Normal |
| Embryo | 8 (16) | 6 | M | | 2.75 | |  | | | n/a | Elective termination | | Normal |
| Embryo | 8 (16) | 6 | F | | 2.5 | |  | | | n/a | Elective termination | | Normal |
| Embryo | 9 (18) | 7 | M | | 5.25 | |  | | | n/a | Elective termination | | Normal |
| Embryo | 9 (18) | 7 | F | | n/k | |  | | | n/a | Elective termination | | Normal |
| Embryo | 10 (21) | 8 | M | | 1.25 | |  | | | n/a | Elective termination | | Normal |
| Embryo | 10 (21) | 8 | F | | n/k | |  | | | n/a | Elective termination | | Normal |
| Embryo | 11 (23) | 9 | M | | n/k | |  | | | n/k | Elective termination | | Normal |
| Embryo | 11 (23) | 9 | F | | 16.67 | | | | | n/k | Elective termination | | Normal |
| Fetus | 12 | 10 | F | | 16.67 | | | | | n/k | Elective termination | | Normal |
| Fetus | 12 | 10 | M | | 16.67 | | | | | n/k | Elective termination | | Normal |
| Fetus | 12 | 10 | n/k | | 3.33 | | | | | n/k | Elective termination | | Normal |
| Fetus | 13 | 11 | M | | 2 | | | | | n/k | Termination of pregnancy | | Normal |
| Fetus | 13 | 11 | M | | n/k | | | | | n/k | Elective termination | | Normal |
| Fetus | 14 | 12 | F | | 2.5 | | | | | n/k | Elective termination | | Normal |
| Fetus | 14 | 12 | F | | 120 |  | | | | 10 | Elective termination | | Normal |
| Fetus | 14 | 12 | F | | 16.67 |  | | | | n/k | Elective termination | | Normal |
| Fetus | 15 | 13 | M | | n/k | | | | | n/k | Termination of pregnancy | | Normal |
| Fetus | 16 | 14 | F | | 16.00 | | | | | n/k | Elective termination | | Normal |
| Fetus | 16 | 14 | M | | n/k | | | | n/k | | Termination of pregnancy | | Normal |
| Fetus | 17 | 15 | M | | n/k | | | | n/k | | Termination of pregnancy | | Normal |
| Fetus | 17 | 15 | F | | n/k | | | | n/k | | Elective termination | | Normal |
| Fetus | 18 | 16 | M | | n/k | | | | 33 | | Elective termination | | Normal |
| Fetus | 18 | 16 | M | | 16.67 | | | | n/k | | Elective termination | | Normal |
| Fetus | 19 | 17 | F | | 144 | | | | 41 | | Elective termination | | Normal |
| Fetus | 19 | 17 | M | | n/k | | | | n/k | | Miscarriage | | Normal |
| Fetus | 20 | 18 | M | 16.67 | | | | | n/k | | Elective termination | |  |
| Fetus | 21 | 19 | M | | 48 | | | | 65 | | Termination of pregnancy | | Normal |
| Fetus | 21 | 19 | M | | 24 | | | | 67 | | Miscarriage | | Normal |
| Fetus | 22 | 20 | M | | 72 | | | | 93 | | Miscarriage | | Normal |
| Fetus | 23 | 21 | F | | n/k | | | | n/k | | Termination of pregnancy | | Normal |
| Fetus | 25 | 23 | M | | 48 | | | 121 | | | Termination of pregnancy | | Normal |
| Fetus | 25 | 23 | M | | n/k | | | 110 | | | Intrauterine death | | Normal |
| Fetus | 26 | 24 | F | | n/k | | | 125 | | | Preterm birth | | Normal |
| Fetus | 27 | 25 | F | | 6 | | | n/k | | | Intrauterine death | Normal | |
| Fetus | 27 | 25 | F | | n/k | | | n/k | | | Preterm birth, defective placenta | Normal | |
| Fetus | 28 | 26 | M | | 96 | | | 159 | | | n/k | Normal | |
| Fetus | 28 | 26 | M | | n/k | | | n/k | | | n/k | Normal | |
| Fetus | 30 | 28 | F | | 240 | | | n/k | | | Maternal road accident/placental separation | Normal frontal histology | |
| Fetus | 34 | 32 | M | | 144 | | | 312 | | | Respiratory distress syndrome | Normal | |
| Fetus | 35 | 33 | F | | n/k | | | 295 | | | Pulmonary hypoplasia, preterm birth | Normal | |
| Fetus | 38 | 36 | F | | 17 | | | 380 | | | Preterm birth | Normal | |
| Fetus | 40 | 38 | M | | 5 | | | n/k | | | Perinatal asphyxia, cardiorespiratory arrest | Normal | |
| Fetus | 40 | 38 | F | | 5 | | | n/k | | | Placental disruption, *in utero* death | Normal | |
| Fetus | 40 | 38 | M | | n/k | | | 370 | | | Septicaemia, premature birth | Normal | |
| Fetus † | 40 | 38 | M | | 72 | | | 356 | | | Intrauterine death | Hypoxic injury | |
| Total number of cases shown is n = 52. In total, 63 cases were assessed. † Only one hypoxic case is included in the table above for reference. Refer also to supplementary figure 1 for a representative example of excluded cases due to hypoxic injury to the area. *BW*: brain weight; *CS:* Carnegie stage; *F:* female; *GA:* gestational age; *M:* male; *n/a:* not applicable; *n/k:* not known; *PCW*: postconceptional week; *PMI*: post-mortem interval. | | | | | | | | | | | | | |
