## Supplemental Table 3 for "Spatiotemporal dynamics of human microglia are linked with brain developmental processes across the lifespan"

| **Supplementary Table 2: Postnatal demographics** | | | | | | | | | | |
| --- | --- | --- | --- | --- | --- | --- | --- | --- | --- | --- |
| **Case** | **Age (years)** | **Sex** | **PMI (h)** | | | **BW (g)** | **Cause of Death** | | | **Histology** |
| Infant | 0.003* | F | | n/k | | 390 | Intrapartum asphyxia | | | Normal |
| Infant | 0.008* | F | | n/k | | 400 | SUDI | | | Normal |
| Infant | 0.01* | M | | n/k | | 415 | Intrapartum asphyxia | | | Normal |
| Infant | 0.02* | M | | n/k | | 375 | n/k | | | Normal |
| Infant | 0.03* | M | | n/k | | 350 | SUDI | | | Normal |
| Infant | 0.03* | F | | n/k | | 335 | SUDI | | | Normal |
| Infant | 0.05* | M | | n/k | | n/k | Respiratory arrest | | | Normal |
| Infant | 0.07* | F | | n/k | | 395 | Intrapartum asphyxia | | | Normal |
| Infant | 0.07* | M | | n/k | | 465 | SUDI | | | Normal |
| Infant | 0.08* | F | | n/k | | 465 | SUDI | | | Normal |
| Infant | 0.08* | F | | n/k | | 525 | SUDI | | | Normal |
| Infant | 0.14* | F | | n/k | | 535 | Pneumonia | | | Normal |
| Infant | 0.17* | M | | n/k | | 530 | SUDI | | | Normal |
| Infant | 0.17* | M | | n/k | | 555 | SUDI | | | Normal |
| Infant | 0.25* | M | | 72 | | 598 | SUDI | | | Normal |
| Infant | 0.33* | M | | n/k | | 840 | SUDI | | | Normal |
| Infant | 0.42* | F | | n/k | | 485 | SUDI | | | Normal |
| Infant | 0.5* | M | | n/k | | 885 | SUDI | | | Normal |
| Infant | 0.50* | M | | n/k | | n/k | n/k | | | Normal |
| Infant | 0.92* | F | | 96 | | 1008 | SUDEP | | | Normal |
| Infant | 0.92* | M | | 96 | | 1288 | Subdural haemorrhage | | | Normal |
| Infant | 1 | M | | 48 | | 1342 | SUDEP | | | Normal |
| Infant | 1.5 | F | | n/k | | 1295 | Aspiration, SUDEP | | | Normal |
| Infant | 2 | M | | 96 | | 1379 | SUDEP | | | Normal |
| Adult | 18 | M | | 5 | | n/k | Car crash |  | | Normal |
| Adult | 25 | M | | 53 |  | 1640 | Suspension by ligature | | | Normal |
| Adult | 25 | M | | 81 |  | 1500 | Road traffic collision | | | Normal |
| Adult | 27 | M | | 67 |  | 1540 | Ischaemic heart disease | | | Normal |
| Adult | 30 | M | | 71 |  | 1670 | n/k | | | Normal |
| Adult | 35 | F | | 44 |  | 1240 | n/k | | | Normal |
| Adult | 37 | F | | 46 |  | 1290 | Hepatic failure | | | Normal |
| Adult | 40 | M | | 48 | | 1320 | Pulmonary embolism | | | Normal |
| Adult | 42 | M | | 103 |  | 1560 | Suspension by ligature | | | Normal |
| Adult | 51 | M | | 51 |  | 1460 | Coronary artery atheroma | | | Normal |
| Adult | 56 | M | | 44 |  | 1500 | Ischaemic heart disease | | | Normal |
| Adult | 53 | M | | 64 |  | 1520 | Depressive episode | | | Normal |
| Adult | 57 | F | | 73 |  | 1320 | Sudden death | | | Normal |
| Adult | 59 | F | | 50 |  | 1500 | Cardiomegaly | | | Normal |
| Adult | 60 | M | | 52 |  | 1460 | Ischaemic heart disease | | | Normal |
| Adult | 66 | M | | 48 |  | 1350 | Hypertensive heart disease | | Normal | |
| Adult | 71 | F | | 41 |  | 1210 | Ischaemic heart disease | | Normal | |
| Adult | 74 | F | | 41 |  | 1520 | Pulmonary embolism | | Normal | |
| Adult | 74 | M | | 23 |  | 1350 | Ischaemic heart disease | | Normal | |
| Adult | 74 | M | | 12 |  | 1600 | Pulmonary embolism | | Normal | |
| Adult | 75 | F | | 24 |  | 1580 | Ischaemic heart disease | | Normal | |
| Total number of cases shown is n = 45, total cases assessed is 71 (Excluded cases due to trauma, malformation, hypoxic injury to the area or loss of antigenicity). *BW:* brain weight; *F:* female; *M:* male; *n/k:* not known; *PMI:* post-mortem interval; *SUDEP*: sudden unexpected death in epilepsy; *SUDI*: sudden unexpected death in infancy; **Age range in days for cases < 1 year*: (1 day - 335 days)*.* | | | | | | | | | | |
